# Redox protein Memo1 coordinates FGF23-driven signaling and small Rho-GTPases in the mouse kidney

**DOI:** 10.1101/2020.12.04.402511

**Authors:** Katalin Bartos, Suresh Krishna Ramakrishnan, Sophie Braga-Lagache, Barbara Hänzi, Fanny Durussel, Arjun Prakash Sridharan, Yao Zhu, David Sheehan, Nancy E. Hynes, Olivier Bonny, Matthias B. Moor

**Author notes:** Co-first authors (Katalin Bartos, Suresh Krishna Ramakrishnan). Correspondence to: Matthias Moor, MD PhD, University Hospital Bern, Department of Nephrology and Hypertension, Freiburgstrasse 15, 3010 Bern, Switzerland.

## Abstract

*Memo1* deletion in mice causes premature aging and an unbalanced metabolism partially resembling *Fgf23* and *Klotho* loss-of-function animals. We report a role for Memo’s redox function in renal FGF23-Klotho signaling using mice with postnatally induced Memo deficiency in the whole body (cKO). Memo cKO mice showed impaired FGF23-driven renal ERK phosphorylation and transcriptional responses. FGF23 actions involved activation of oxidation-sensitive protein phosphotyrosyl phosphatases (PTP) in the kidney. Redox proteomics revealed excessive thiols of Rho-GDP dissociation inhibitor 1 (Rho-GDI1) in Memo cKO, and we detected a functional interaction between Memo’s redox function and oxidation at Rho-GDI1 Cys79. In isolated cellular systems, Rho-GDI1 did not directly affect FGF23-driven cell signaling, but we detected disturbed Rho-GDI1 dependent small Rho-GTPase protein abundance and activity in the kidney of Memo cKO mice. Collectively, this study reveals previously unknown layers in the regulation of renal FGF23 signaling and connects Memo with the network of small Rho-GTPases.

## Introduction

Fibroblast growth factor (FGF) 23 plays a critical role during the progression of chronic kidney disease (CKD). Elevated FGF23 concentration is associated with complications such as vascular calcification, heart disease, and mortality in patients with and without CKD (Marthi et al. 2018). FGF23 is secreted by osteoblasts or osteocytes upon stimuli such as vitamin D, parathyroid hormone, extracellular phosphate, or inflammatory cytokines (Richter and Faul 2018). FGF23 activates a canonical pathway of FGFR1 using Klotho as a co-receptor to stimulate ERK signaling in parathyroid glands or kidneys (Urakawa et al. 2006). Aberrant renal FGF23-driven signaling in experimental models of CKD has detrimental cellular consequences such as the induction of inflammatory responses (Dai et al. 2012), impaired leukocyte recruitment mediated by FGFR2 (Rossaint et al. 2016), and renal fibrosis (Smith et al. 2017; Hao et al. 2021). However, the underlying molecular mechanisms are not fully understood.

Mediator of Cell Motility 1 (Memo) is an evolutionarily conserved protein associated with the cytoskeleton and with receptor tyrosine kinases (Marone et al. 2004; Zaoui et al. 2008; Jiang et al. 2013; Schotanus and Van Otterloo 2020). Memo structurally resembles bacterial nonheme iron dioxygenases (Qiu et al. 2008) and has a copper-reducing redox function. Memo is required for activity of nicotinamide adenine dinucleotide phosphate (NADPH) oxidase (NOX) in mammalian cell protrusions (MacDonald et al. 2014). In mice, *Memo1* deletion resulted in embryonic lethality. The postnatally-inducible ubiquitous deletion of *Memo1* exon 2 (Memo cKO) using Cre-lox techniques resulted in a mouse phenotype of premature aging with dwarfism, alopecia, hair graying, loss of subcutaneous fat, hypogonadism, and abnormal gait (Haenzi et al. 2014). Moreover, these animals showed a hypersensitivity to insulin and an increased glucose tolerance compared to controls (Haenzi et al. 2014). Such findings are rarely seen in models of premature aging and strikingly resemble the widely studied *Fgf23* and *Klotho*-deficient mouse models (Kuro-o et al. 1997; Shimada et al. 2004). Moreover, four to eight weeks after undergoing *Memo1* exon 2 excision recombination, Memo cKO animals developed kidney failure, elevated serum concentrations of FGF23 and calcium, and a bone and mineral disease (Haenzi et al. 2014; Moor et al. 2018; Moor et al. 2020). This partially overlapped with *Fgf23* and *Klotho*-deficient strains (Kuro-o et al. 1997; Shimada et al. 2004). However, the hyperphosphatemia of *Fgf23* and *Klotho* loss-of-function strains (Kuro-o et al. 1997; Shimada et al. 2004) was not found in Memo cKO mice. Memo cKO mice, however, also present increased renal epidermal growth factor expression and increased levels of magnesium transport proteins and serum concentrations (Haenzi et al. 2014; Moor et al. 2020), which are tightly linked to the function of the distal nephron (Groenestege et al. 2007). Epidermal growth factor signaling promotes renal phosphate excretion, and this may explain the normal phosphaturia in Memo cKO (Arar et al. 1995; Arar et al. 1999).

Results from cell culture experiments nevertheless support the hypothesis that FGF23, the FGFR co-receptor klotho, and Memo act in a common signaling pathway: First, Memo co-immunoprecipitated with FGFR1 and with FRS2 and further adaptor proteins that are recruited to the FGFR upon ligand binding. Second, Memo knockdown in cells diminished the phosphorylation of FRS2 induced FGF2 or FGF23 treatment (Haenzi et al. 2014).

Whether these findings translate to a complex organ such as the kidney, the main site of FGF23 action, is unclear. Moreover, the mechanisms how Memo affects RTK signaling beyond a potential role as a scaffold for adaptor proteins (Marone et al. 2004; Haenzi et al. 2014; Newkirk et al. 2018) are incompletely understood.

Here, we hypothesized that Memo modulates renal FGF23-induced cell signaling by its function as a redox enzyme. In the experiments presented here, we demonstrate that Memo has a pivotal role in FGF23-driven signaling in the kidney. We uncover that Memo regulates activity and abundance of small Rho-GTPases and protein cysteine oxidation that accompanies intracellular signaling responses to FGF23.

## Materials and Methods

### Animal studies

Mice were fed standard chow (TS3242 Kliba Nafnag, Kaiseraugst, Switzerland) and were kept on 12/12 or 14/10 light-dark cycles. Mice floxed for exon 2 of the *Memo1* gene (Haenzi et al. 2014) backcrossed to C57BL/6J background over 10 generations were crossed with CreERTM transgenic mice (Hayashi and McMahon 2002) carrying a tamoxifen-inducible Cre recombinase controlled by a beta-actin promoter/enhancer. Genotypes were determined by PCR of ear punch biopsy DNA using primers: Memo forward 5’-*CCCTCTCATCTGGCTTGGTA*-3’, Memo reverse 5’-*GCTGCATATGCTCACAAAGG*-3’, Cre forward 5’-*AGGTTCGTGCACTCATGGA*-3’, Cre reverse 5’-*TCACCAGTTTAGTTACCC*-3’. For the current study, all animals used were males, with the exception of the experiments reported in Figure 5 in which there was a sex ratio of 50:50 in each genotype. Loss of Memo was induced by 3 daily intraperitoneal injections with 2mg tamoxifen at age 4 weeks (T5648 Sigma-Aldrich, distributed through Merck, Buchs Switzerland). Memo^fl/fl^ littermates without Cre but treated with tamoxifen served as controls. For FGF23 treatments, mice aged 6.5 weeks were starved for 6 hours and intraperitoneally injected with 220ng/g body weight of recombinant mouse FGF23 (2629-FG-025 R&D Systems, Minneapolis, MN, USA) in PBS-BSA 0.1% or PBS-BSA 0.1% vehicle with injection volume of 4μL/g body weight. One hour later mice were dissected, a protocol adapted from (Gattineni et al. 2014). Treatments were randomly allocated by flipping a coin. Animals were euthanized after 1h by terminal exsanguination under anesthesia. For the indicated pre-treatments to modulate FGF23 signaling, animals additionally received intraperitoneal injections of with sodium orthovanadate 20mg/kg (13721-39-6 Sigma-Aldrich) (Wang et al. 2013) or sodium chloride 0.9% 1h before their injection of FGF23 or PBS-BSA 0.1%.

### Cell culture and transfection

HEK293 cells were obtained from ATCC and were tested negative for mycoplasma using LookOut® PCR-based kit (MP0035-1KT Sigma-Aldrich). HEK293-Klotho cells stably expressing Klotho were a gift from Dr. Bettina Lorenz-Depiereux, HelmholtzZentrum München and were described in (Diener et al. 2015). Cells were transfected with Myc-DDK-tagged Rho-GDI1 (ARHGDIA) (MR202112 OriGene) using Lipofectamine 3000 (L3000015 Invitrogen, distributed through Thermo Fisher Scientific, Reinach, Switzerland). HEK293-Klotho cells were seeded at density of 500 000 cells/well on 6-well tissue culture plates and transfected with RhoGDI1 (ARHGDIA) (NM_004309) Human Tagged ORF Clone (RG200902 OriGene) using Lipofectamine 3000 Transfection Reagent (L3000015 Invitrogen, distributed through Thermo Fisher Scientific, Reinach, Switzerland) following the manufacturer’s protocol. 72 hours after transfection, cells were treated with vehicle (0.1% BSA) or with 100ng/ml FGF23 (SRP3039 Sigma-Aldrich, Buchs, Switzerland) for 5 or 15 minutes. Following treatment, cellular protein was extracted in RIPA (R0278 Sigma-Aldrich) by 1 hour incubation on a shaker, followed by centrifugation at 10000g.

### Cellular siRNA experiments

Hek293-Klotho cells were seeded at density of 200 000 cells/well on 12-well tissue culture plates and transfected with ARHGDIA Ambion™ Silencer™ Select Pre-Designed siRNA (4427038 Ambion, distributed through Thermo Fisher Scientific, Reinach, Switzerland) or with RhoGDI1 (ARHGDIA) Human siRNA Oligo Duplex (Locus ID 396) (SR300287 OriGene Technologies Inc., Rockville MD, USA) using Lipofectamine 3000 Transfection Reagent (L3000015 Invitrogen, distributed through Thermo Fisher Scientific, Reinach, Switzerland) following the manufacturer’s protocol. 72 hours after transfection with Rho-GDI1 or negative control siRNAs, cells were treated with vehicle (0.1% BSA) or with 100ng/ml FGF23 (SRP3039 Sigma-Aldrich, Buchs, Switzerland) for 5 or 15 minutes. Following treatment, cellular protein was extracted in RIPA (R0278 Sigma-Aldrich) by 1 hour incubation on a shaker, followed by centrifugation at 10000g.

### Redox proteomics

Kidney halves pooled from 2 mice per sample were homogenized in 10 mM Tris-HCl, pH 7.2, 250 mM sucrose, 1 mM EDTA, 150 mM KCl and 1 mM PMSF and spun down. Protein thiols and carbonyls were labelled with either 0.2 mM 5’-iodoacetamido fluorescein (IAF) or 1mM fluorescein-5-thiosemicarbazide (FTSC), respectively, and incubated for 150 min at 37 °C in the dark. Proteins were precipitated with 20 % TCA and centrifuged at 20,000 ×g (3 min, 4 °C). Protein pellets were resuspended and washed with 100% ethanol/ethyl acetate (1:1) and 96% acetone, respectively, for carbonyl and thiol groups. Pellets were resuspended in Tris-HCl 0.5 M pH 6.8, glycerol 10%, SDS 0.5% and bromophenol blue and applied to SDS-PAGE. Gels were scanned for fluorescence in a Typhoon Trio Scanner 9400 (Control v5.0 + variable Mode Imager-RA 501: PRT<I/06/004, GE Healthcare, Buckingshamshire, UK; excitation, 490-495 nm; emission, 515–520 nm). Protein-associated fluorescence intensities (arbitrary units, AU) were analyzed using Quantity One image analysis software (BioRad, Hercules, CA, USA). Gels were stained with Colloidal Coomassie Brilliant Blue G250 and scanned.

For 2-dimensional analysis by gel electrophoresis (2DGE), proteins were first separated by pI (first dimension: isoelectric focusing IEF), followed by orthogonal separation according to molecular weight (second dimension: SDS-PAGE). Proteins were rehydrated in 5 M urea, 2 M thiourea, 2% CHAPS, 4% ampholyte (Pharmalyte 3-10, Amersham-Pharmacia Biotech, Little Chalfont, Bucks, UK), 1% Destreak reagent (Amersham-Pharmacia Biotech, Buckinghamshire, UK), and trace amounts of bromophenol blue, and then immobilized in 7 cm IPG strips (pH 3-10 : 70×3×0.5 mm) and a linear gradient (NL) (GE Healthcare Immobiline™ Dry Strip IPG, GE17-6001-11 Bio-Sciences AB, Bio-Rad, Hercules, CA, USA). Proteins were focused at room temperature in a Protean IEF Cell (Bio-Rad) for at least 15 h, according to the following steps: (1) a linear voltage increase until 250 V for 15 min, (2) 10,000 V for 2 h (50 μA/strip), (3) focusing at 20,000 V, and (4) hold at 500 V. Following IEF, strips were equilibrated for 20 min in equilibration buffer (6 M urea, 0.375 M Tris, pH 8.8, 2% SDS, 20% glycerol, containing 2% DTT) and then for 20 min in equilibration buffer containing 2.5% iodoacetamide. IPG strips were loaded onto 12% SDS-PAGE gels (PROTEAN Plus Dodeca Cell, BioRad). Gels were scanned for fluorescence as above and stained with Coomassie Blue followed by densitometry. Progenesis SameSpots Software (S/No.62605/3787; Nonlinear USA Inc, Durham, NC USA) was used to normalize FTSC/ IAF-labeled protein spots and Coomassie-staining intensity. Fluorescence spots were normalized to protein intensity for the same gel.

All experiments were performed in triplicates. Images of 2D gels were subjected to landmarking alignment so that corresponding spots were matched with each other, based on 3D Gaussian distribution after raw image correction and background subtraction. Spot intensities were normalized. Differences between protein spots in 2D gel images were automatically determined.

Spots with differences of p<0.05 by ANOVA were manually excised from colloidal Coomassie-stained 2-DE gels. Proteins were extracted and enzymatically digested using a Perkin Elmer – Janus automated workstation. Following digestion, samples were reconstituted in 0.1% formic acid and analyzed using a Dionex U3000 Liquid Chromatography System (Dionex, Sunnyvale, CA, USA) and the Daltonics HCT Ion Trap Mass Spectrometer (Bruker, Glasgow, UK). The peptide fragment mass spectra were acquired in data-dependent AutoMS(2) mode with a scan range of 300–1500 m/z, and up to three precursor ions were selected from the MS scan (100–2200 m/z). Precursors were actively excluded within a 1.0-min window, and all singly charged ions were excluded. Following LC-MS/MS, Mascot Generic Files were created and the MASCOT database NCBInr was searched using Matrix Science webserver (www.matrixscience.com). The default search parameters were: enzyme = trypsin; maximum number of missed cleavages = 1; fixed modifications = carbamidomethyl (C); variable modifications = oxidation (M); peptide tolerance ±1.5 Da; MS/MS tolerance ± 0.5 Da; peptide charge = 2+ and 3+. Identified proteins were considered if a MASCOT score above 95% confidence was obtained (p < 0.05) and at least one peptide was identified with a score above 95% confidence (p < 0.05). This analysis was conducted at the Proteomics Core Facility of the University of Aberdeen, UK. All mass spectrometry proteomics data from these experiments have been deposited to the ProteomeXchange Consortium via the MassIVE repository with the dataset identifier PXD022342.

### Cysteine oxidation analyses of recombinant protein

Amino acids 24-204 of human Rho-GDI1/ARHGDIA protein expressed in *E. coli* was obtained from Novusbio (NBP1-50861, Centennial, CO, USA NBP1-50861), with >95% purity by SDS-PAGE and supplied in 20mM Tris-HCl buffer (pH 8.0), 1mM DTT, 10% glycerol without preservatives. Recombinant human Memo expressed in *E. coli* with >95% purity, supplied in 20 mM Tris-HCl buffer, pH 8.0, 50% glycerol, 5 mM DTT, 300 mM NaCl, 2 mM EDTA was from Antibodies-online (ABIN2130536, Aachen, Germany). Copper-reducing oxidant activity of the batch of recombinant Memo protein used in this experiment was determined and reported in (Moor et al. 2020).

First, as a putative metal cofactor pre-loading step, 2μg Memo was incubated with RCM buffer with or without 10μM CuCl_2_. After 30 minutes incubation at 4°C the free CuCl_2_ was removed by Slide-A-Lyzer™ MINI Dialysis Device, 3.5K MWCO (69550 Thermo Fisher Scientific) against RCM buffer for 30 minutes at 4°C. Proteins were incubated for 15 minutes as 1000ng Rho-GDI1 alone; 1000ng Rho-GDI1 with 250ng Memo; 1000ng Rho-GDI1 with 250ng CuCl_2_ pretreated Memo; and 1000ng Rho-GDI1 with 100μM of hydrogen peroxide as a positive control (H1009 Sigma-Aldrich), all in duplicates for each experiment, to individually label the total cysteine and the oxidized cysteine content of Rho-GDI1.

The samples for total cysteine content labeling were reduced with 5mM Tris (2-carboxyethyl) phosphine (TCEP) (646547 Sigma-Aldrich). Total cysteine content was labeled with tandem mass tag (TMT) iodoTMTsixplex™ (90102 Thermo Fisher Scientific) followed by quenching with 20mM 1DTT (43816 Sigma-Aldrich).

The samples for oxidized cysteine content labeling were treated with 500mM iodoacetamide (I6125 Sigma-Aldrich) for 30 minutes followed by quenching with 20mM DTT and reducing with 5mM TCEP. The oxidized cysteine content of the samples was labeled with iodoTMTsixplex™.

Samples were precipitated with the SDS-PAGE Clean-Up Kit (10074304 GE Healthcare Life Sciences, distributed through Thermo Fisher Scientific) and separated on SDS-PAGE, followed by Coomassie Blue staining and manual band excision. Proteins were digested by chymotrypsin at 50°C for 5 hours and analyzed by LC-MS/MS (PROXEON coupled to a QExactive HF mass spectrometer, Thermo Fisher Scientific) with two injections of 5μl digests. Peptides were trapped on a μPrecolumn C18 PepMap100 (5μm, 100 Å, 300 μm×5mm, Thermo Fisher Scientific) and separated by backflush on a C18 column (5μm, 100 Å, 75 μm×15 cm, C18) by applying a 20-minute gradient of 5% acetonitrile to 40% in water, 0.1% formic acid, at a flow rate of 350nl/min. The Full Scan method was set with resolution at 60,000 with an automatic gain control (AGC) target of 1E06 and maximum ion injection time of 50ms. The data-dependent method for precursor ion fragmentation was applied with the following settings: resolution 15,000, AGC of 1E05, maximum ion time of 110 milliseconds, mass window 1.6*m*/*z*, first mass 100m/z, collision energy 27, under fill ratio 1%, charge exclusion of unassigned and 1+ ions, and peptide match preferred, respectively. Spectra were interpreted with Proteome Discoverer 2.4.0.305, with chymotrypsin rules allowing up to 8 missed cleavages, using variable modification of carboamidomethylated (+57.021Da), dioxidation (+31.990Da), trioxidation (+47.985Da) and iodoTMT labelling (+324.216Da) on Cys, and variable modification of oxidation (+15.99Da) on Met, and acetylation (+42.011Da) on protein N-Term. Parent and fragment mass tolerances were set to 10ppm and 0.02Da, respectively. Strict target false discovery rate for highly confident peptide-spectrum matches was set to 0.01. Protein identifications were only accepted when two unique peptides fulfilling the 1% FDR criterium were identified. This analysis was performed at the Proteomics Mass Spectrometry Core Facility of University of Bern. Resulting intensity peaks of specific peptide modifications were normalized by total signal intensity and are displayed as heatmap of non-transformed z-scores, i.e. a subtraction of the row mean followed by a division by the row standard deviation. All mass spectrometry proteomics data from recombinant protein analyses have been deposited to the ProteomeXchange Consortium via the PRIDE repository with the dataset identifier PXD022382.

### Western blot

For immunoblotting, lysates of cells and tissues were lyzed in RIPA buffer or NP-40 buffer (150mM NaCl, 50 mM HEPES pH7.4, 25mM NaF, 5mM EGTA, 1mM EDTA, 1% Nonidet P-40, 2M Na ortho-vandate and 1mM DTT supplied with protease inhibitors leupeptin 10μg/L, aprotinin 10μg/L, and PMSF 1mM) were prepared and denatured. Proteins were separated by SDS-PAGE and transferred onto nitrocellulose or PVDF, stained by Ponceau S, blocked in dried nonfat milk 5%-TBST or bovine serum albumin (A9647 Sigma-Aldrich) 3% in TBST before incubation with primary antibodies against Memo (1:2000, produced in-house (Haenzi et al. 2014) or 1:1000, HPA042603 Sigma-Aldrich), pERK (1:1000, sc-7383 Santa Cruz, Dallas TX, USA), tERK (1:1000, sc-93 Santa Cruz), actin (1:2000, A2066 Sigma-Aldrich), Rho-GDI1 (1:500, ABIN969501 Antibodies-online), Rac1 (1:500, ab33186 Abcam, Cambridge UK), RhoA (1:500, NBP2-22529 NovusBio) and Klotho (1:1000, AF1819 R&D Systems). Membranes were incubated with anti-mouse or anti-rabbit horseradish peroxidase-conjugated secondary antibodies (Milian Analytica, Rheinfelden Switzerland or ImmunoResearch, distributed through LubioScience GmbH, Zürich, Switzerland) and imaged using Fusion Solo (Witec AG, Sursee Switzerland) or a ChemiDoc XRS+ System (BioRad). For reprobing, membranes were stripped using a low pH buffer (25mM glycine-HCl, pH2, 0.4% (w/v) SDS).

### Biochemical analyses

Serum electrolytes were analyzed by the Lausanne University Hospital: Total calcium (NM-BAPTA method), phosphate (phosphomolybdate method), and creatinine (modified Jaffé method). Intact FGF23 was analyzed using an ELISA (Kainos Japan CY-4000) following the manufacturer’s instructions. Protein phosphotyrosyl phosphatase (PTP) activity was measured in kidney lysates using fluorometric Protein Tyrosine Phosphatase Activity Assay Kit (#K829, BioVision Inc., Milpitas CA, USA) according to the manufacturer’s instructions. For quantification of Rho-GTPase activity in kidney homogenates, colorimetric Rac1 and RhoA G-LISA Activation Assay Kits (BK128-S and BK124-S Cytoskeleton Inc., Denver CO, USA) were used according to the manufacturer’s instructions.

### RNA isolation

Frozen kidneys were homogenized with metal beads in a TissueLyser (Qiagen, Hombrechtikon, Switzerland). RNA was extracted using TRI Reagent Solution (AM9738 Ambion, Austin, TX, USA) for all downstream applications.

### qPCR

One μg of RNA per kidney half was reverse-transcribed using PrimeScript RT Reagent kit (RR037 TAKARA, Shiga, Japan). Two μl of cDNA was used for quantitative real-time PCR to assess gene mRNA expression. Assays were performed using SYBR Green (Applied Biosystems, Foster City, CA UA) on a 7500 Fast machine (Applied Biosystems). Samples were run in triplicates in 20uL total volume for each gene, and actin or GAPDH was used for normalization. Melting curves were obtained for every run. Program settings were: 95°C 20 seconds, 40 cycles (95°C 3 sec, 60°C 30 sec), and for melting curve stage: 95°C 15sec, 60°C 1min, rising at 1% ramp speed to 95°C (15sec), 60°C 15sec. Data were analyzed using the delta-delta CT method. Primers were ordered from Microsynth (Balgach, Switzerland), and sequences are shown in Supplemental table 1.

**Table 1:** Identification of altered proteins in redox proteomics of the kidney.

|  | Spot no. | Identified protein | Mw | Protein score | Total ion score | coverage (%) | N of peptides identified | P Value | Average normalized volumes |  |
| --- | --- | --- | --- | --- | --- | --- | --- | --- | --- | --- |
|  |  |  |  |  |  |  |  |  | Ctrl-V | Memo cKO-V |
| Coomassie | 386 | 60 kDa heat shock protein, mitochondrial | 61 | 666 | 447 | 36 | 19 | 0.008 | 1.68E+05 | 2.69E+05 |
|  | 360 | Succinyl-CoA:3-ketoacid coenzyme A transferase 1, mitochondrial | 56 | 360 | 299 | 26 | 10 | 3.47E-05 | 2.67E+05 | 1.53E+05 |
|  | 416 | Alpha-1-antitrypsin 1-5 | 46 | 54 | 34 | 13 | 3 | 0.008 | 6.95E+04 | 3.41E+04 |
|  | 628 | Phosphatidylethanolamine-binding protein 1 | 21 | 131 | 110 | 25 | 2 | 0.038 | 4.00E+05 | 1.86E+05 |
|  | 534 | Aminoacylase-1 | 46 | 126 | 107 | 14 | 7 | 0.033 | 2.27E+05 | 1.27E+05 |
| Thiols | 175 | Actin, cytoplasmic 1 | 42 | 375 | 246 | 32 | 13 | 0.047 | 2.49E+07 | 1.23E+07 |
|  | 78 | Heat shock cognate 71 kDa protein | 71 | 433 | 250 | 22 | 10 | 0.024 | 3.39E+06 | 2.03E+06 |
|  | 244 | Pyruvate dehydrogenase E1 component subunit beta, mitochondrial | 39 | 374 | 193 | 37 | 13 | 9.58E-04 | 7.69E+05 | 3.30E+06 |
|  | 132 | Glutathione synthetase | 52 | 684 | 495 | 38 | 16 | 0.035 | 3.01E+06 | 1.28E+06 |
|  | 202 | L-lactate dehydrogenase B chain | 37 | 611 | 334 | 52 | 19 | 0.001 | 6.05E+06 | 3.18E+06 |
|  | 216 | Malate dehydrogenase, cytoplasmic | 37 | 321 | 296 | 31 | 10 | 0.024 | 1.02E+07 | 3.83E+06 |
| Carbonyls | 50 | 78 kDa glucose-regulated protein | 72 | 641 | 415 | 28 | 15 | 0.047 | 2.33E+06 | 8.25E+05 |
|  | 32 | Serotransferrin | 79 | 769 | 484 | 36 | 20 | 0.028 | 1.20E+06 | 2.72E+06 |
|  | 117 | Protein disulfide-isomerase A3 | 57 | 487 | 286 | 32 | 13 | 0.015 | 4.22E+05 | 1.94E+05 |
|  |  |  |  |  |  |  |  |  | Ctrl-F | Memo cKO-F |
| Coomassie | 535 | Aminoacylase-1 | 46 | 231 | 137 | 21 | 7 | 0.008 | 5.93E+05 | 3.46E+05 |
|  | 507 | Alpha-enolase | 47 | 414 | 285 | 23 | 7 | 0.02 | 1.69E+05 | 3.07E+05 |
|  | 735 | Periaxin | 148 | 36 | 28 | 6 | 10 | 0.013 | 4.32E+05 | 7.26E+05 |
| Thiols | 236 | Selenium-binding protein 1 | 53 | 997 | 665 | 52 | 29 | 0.024 | 3.06E+06 | 6.32E+06 |
|  | 267 | Malate dehydrogenase, cytoplasmic | 37 | 552 | 506 | 46 | 20 | 0.001 | 5.16E+05 | 3.19E+06 |
|  | 378 | Rho GDP-dissociation inhibitor 1 | 23 | 116 | 92 | 20 | 2 | 0.035 | 1.83E+06 | 3.10E+06 |
|  | 212 | Glutathione synthetase | 52 | 185 | 106 | 28 | 8 | 0.008 | 6.58E+06 | 1.70E+06 |
|  | 253 | L-lactate dehydrogenase B chain | 37 | 575 | 321 | 48 | 15 | 0.04 | 1.93E+06 | 1.18E+06 |
|  | 305 | Pyruvate dehydrogenase E1 component subunit beta, mitochondrial | 39 | 366 | 237 | 25 | 7 | 0.009 | 5.09E+06 | 8.79E+05 |
|  | 175 | Actin, cytoplasmic 1 | 42 | 142 | 92 | 12 | 5 | 0.033 | 1.53E+07 | 3.94E+06 |
| Carbonyls | 67 | ATP synthase subunit beta, mitochondrial | 56 | 554 | 356 | 31 | 13 | 0.232 | 6.03E+06 | 1.11E+07 |
|  | 146 | Malate dehydrogenase, cytoplasmic | 37 | 399 | 432 | 30 | 13 | 0.26 | 5.08E+05 | 9.52E+05 |
|  | 96 | Selenium-binding protein 1 | 53 | 691 | 394 | 46 | 22 | 0.156 | 1.19E+06 | 6.80E+05 |

Mw, molecular weight in kDa. Ctrl, control. V, vehicle. F, FGF23. P-Values determined by t-test.

### RNAseq

For RNAseq, extracted RNA from mouse kidney halves was pairwise pooled to 3 individual pools for each of 4 experimental conditions (2 treatment states, 2 genotypes). Quality of mRNA was assessed and verified using a Fragment Analyzer (Advanced Analytical Technologies Inc.). Following quality validation, TruSeq Stranded RNA Sample Prep Kit (20020596 Illumina, San Diego CA, USA) was used with 1ug of total RNA for the construction of sequencing libraries. cDNA was sequenced using the Illumina HiSeq2500 platform and single-end reads (1×100). All sequencing data are accessible on National Center for Biotechnology Information, Sequence Reads Archive, Accession: PRJNA672305. Purity-filtered reads were trimmed with Cutadapt (v. 1.3, Martin 2011), filtered using seq_crumbs (v. 0.1.8) and aligned against the *Mus musculus* (Ensembl version GRCh38.82) transcriptome using STAR (v. 2.4.0f1). Data were normalized with the Trimmed Means of M-Values (TMM) method of the edgeR, and transformed with voom method of bioconductor package limma. Genes expressed at low levels were removed when none of the samples had more than 1 count per million reads. Only genes labeled as “protein coding” were retained for the statistical tests. A heatmap was constructed using the heatmap2 function in R 4.1.0 package gplots using z-scores of log-transformed normalized reads.

### Statistical analyses

Analyses of parametric continuous variables were performed by t-test or by ANOVA, followed by post-tests using Bonferroni correction for multiple testing across all experimental groups. Analyses of nonparametric continuous variables were assessed by Kruskal Wallis test, followed by Dunn’s correction for multiple testing across all experimental groups. Analyses were performed in GraphPad Prism 5.0, and two-tailed p-values of <0.05 were considered significant.

Analyses of RNAseq data were performed with limma and DESeq2 (R version 3.2.1, limma version 3.22.7 and DESeq2 version 1.8.1). Because the experiment has a 2×2 factorial design with two mouse genotypes and two treatment states, linear models were generated with all 4 groups as factors. Subsequently, contrasts of interest were extracted using moderated t-tests for the following groups: 1. “KOFGF-KOV” (treatment effect in Memo cKO), 2. “WTFGF-WTV” (treatment effect in floxed control genotype), 3. “KOFGF-WTFGF” (KO genotype effect in FGF23-treated mice), 4. “KOV-WTV” (KO genotype effect in vehicle-treated mice), and 5. “(KOFGF-KOV)-(WTFGF-WTV)” for the interaction between genotype and treatment. P-values were adjusted by the Benjamini-Hochberg method, controlling for false discovery rate (FDR) across all 5 contrasts as a correction of multiple testing conditions. All sequencing data are available on National Center for Biotechnology Information, Sequencing Reads Archive under the identifier PRJNA672305. Gene ontology enrichment was assessed using GOrilla, accessed on Nov 17^th^ 2020 (Eden et al. 2009) using entire genes lists ranked by adjusted p-values.

### Study approval

The present study was approved by the veterinary services of the Cantons of Vaud and Bern, Switzerland.

## Results

### FGF23-driven renal ERK phosphorylation and transcriptional responses depend on Memo

For the present study, we set out to test if FGF23-driven signaling depends on Memo not only in cultured cells, but also in the more complex realm of the rodent. To this end, we generated inducible whole-body Memo cKO, and floxed control mice not harboring the Cre transgene, and maintained them on C57BL/6 background as previously described (Moor et al. 2018). We treated mice of both genotypes with tamoxifen to excise *Memo1* exon 2 in the whole body. We randomized them to intraperitoneal FGF23 or vehicle injections as depicted in (Figure 1A). FGF23 injections yielded a 14-fold increase in circulating intact FGF23 (Supplemental Figure 1A). Previous studies have found deficits in several signaling responses in cultured cells that were Memo-deficient, including ERK signaling (Frei et al. 2016; Schotanus and Van Otterloo 2020). We assessed phosphorylated and total ERK protein levels in whole kidney lysates and detected an increase in phospho-ERK in control mice treated with FGF23, but not in FGF23-treated Memo cKO animals (Figure 1B-C). Next, we analyzed expression of ERK downstream genes in kidney of mice in both genotype and treatment groups. A 1 hour treatment with FGF23 consistently increased *Cyp24a1* (Figure 1D) and *Egr1* expression (Figure 1E) in control mice only, but not in FGF23-treated cKO. These data highlight a role for Memo in promoting FGF23-driven cell signaling in the kidney.

**Figure 1.**
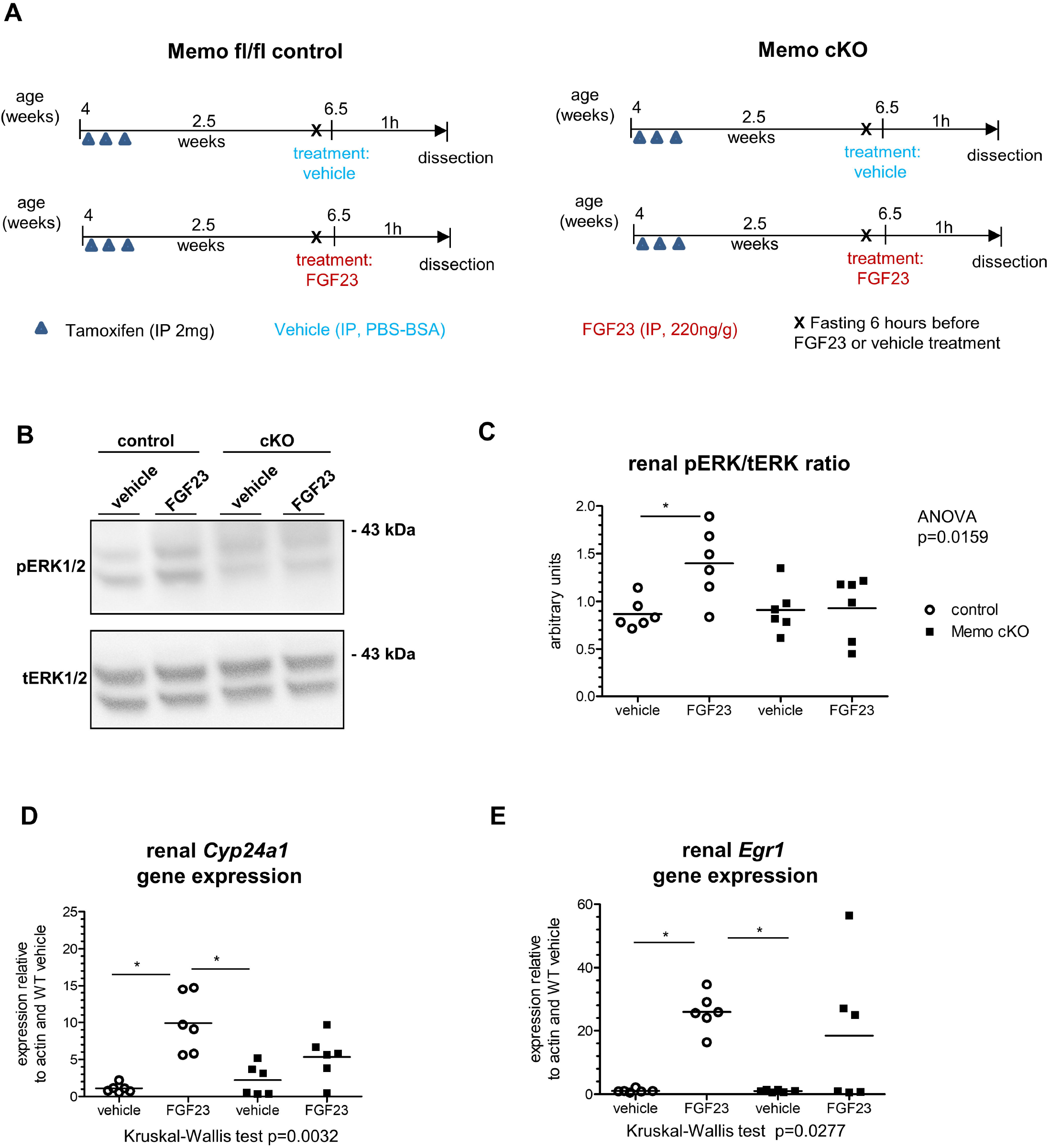
Impaired FGF23-induced renal signaling in whole-body Memo KO (cKO) mice. Mice were randomized to recombinant FGF23 or vehicle intraperitoneal (i.p.) injection 2.5 weeks after Memo ablation (A). Phosphorylated (p) versus total (t) renal ERK protein assessed by Western blot 1 hour after FGF23 or vehicle (B). Densitometric quantification showed a blunted increase in pERK/tERK ratio in response to FGF23 in kidney of Memo cKO (C). Target gene responses of Cyp24a1 (D) and egr1 (E) were assessed by qPCR. Statistical analysis was obtained by ANOVA with Bonferroni’s post-tests (C) or Kruskal-Wallis test with Dunn’s post-test (D-E) between all groups. p *, p<0.05 in post-tests. n=6 per condition (B-E).

These findings were not biased by the severe phenotype of the Memo cKO animals: The mice included in the present study were all studied prior to manifestation of premature aging signs reported previously (Haenzi et al. 2014) and were still thriving as shown by body weight (Supplemental Figure 1B), serum creatinine (Supplemental Figure 1C), calcium and phosphate concentrations (Supplemental Figures 2D-E) that were overall comparable between genotypes. In addition, we ensured for that the KO of Memo was successful in all analyzed samples. (Supplemental Figure 1F). Overall, the mouse model showed Memo depletion as expected, but the observed signaling phenotype was not caused by the disease traits that develop later in this model.

### FGF23-driven changes in transcriptional signatures related to MAPK phosphatase activity depend on Memo

FGFR signaling is controlled by negative regulators. We therefore measured SEF, encoded by *Il17rd*, by qPCR in the kidney of Memo cKO and controls stimulated with FGF23 or vehicle, and *Il17rd* transcripts were found to be unchanged across treatment and genotypes in the kidney (Supplemental Figure 1G). We have previously shown that Memo cKO animals develop kidney failure at age 12-14 weeks or five weeks after recombination, and during advanced CKD, a severe loss of Klotho protein aggravates the inability of the kidney to excrete phosphate (Koh et al. 2001; Moor et al. 2018). We therefore examined protein levels of Klotho in control and KO animals studied at age 6.5 weeks or 2.5 weeks after recombination in the present study. The renal protein quantity of the FGF23 co-receptor Klotho was not changed between the different experimental groups (Supplemental Figure 1H). To summarize, abundance of the co-receptor Klotho or an immediate negative feedback regulator were not altered and therefore unlikely to be responsible for the renal FGF23 signaling phenotype observed in Memo cKO animals.

To gain further insight into FGF23-dependent signaling pathways disturbed in Memo cKO, we assessed the transcriptome of pair-wise pooled kidney samples from FGF23-treated and vehicle-treated Memo cKO and control mice (Figure 2A, Supplemental tables 2-6). In the acute response after 1 hour of FGF23 treatment, we found a consistent genotype effect but no strong treatment effect on the overall transcriptome in non-supervised clustering (Supplemental figure 2A), principal component analyses (Supplemental figure 2B) and in heatmap analysis of log-transformed normalized reads (Figure 2B). Nevertheless, thirteen transcripts were increased by FGF23 in control genotypes and 1 in Memo cKO (Figure 2C, Supplemental figure 2C). Among the 13 FGF23-dependently increased transcripts, 4 encoded dual-specificity phosphatases (DUSP). Consequently, transcripts with a gene ontology functionally annotated to mitogen-activated protein kinase (MAPK or ERK) phosphatase activity were strongly enriched by FGF23 treatment in control mice (Supplemental figure 3, Supplemental table 3). By contrast, FGF23 treatment in Memo cKO yielded a less strong enrichment related to MAPK phosphatase activity (Supplemental figure 4, Supplemental table 2). In an interaction analysis of genotype and FGF23 treatment this effect remained significant (Supplemental figure 5, Supplemental table 6), indicating stronger responses to FGF23 in Memo controls on global transcriptomic levels. Additional analyses of selected transcripts hinted at impaired FGF23-driven changes in *Cyp24a1, Cyp27b1* or *Hbegf*, along with consistently lower Memo1 transcript levels in Memo cKO (Figure 2C-D). Overall, we found a genotype-different renal transcriptional signature in response to FGF23 that involved transcripts related to MAPK phosphatase activity.

**Figure 2.**
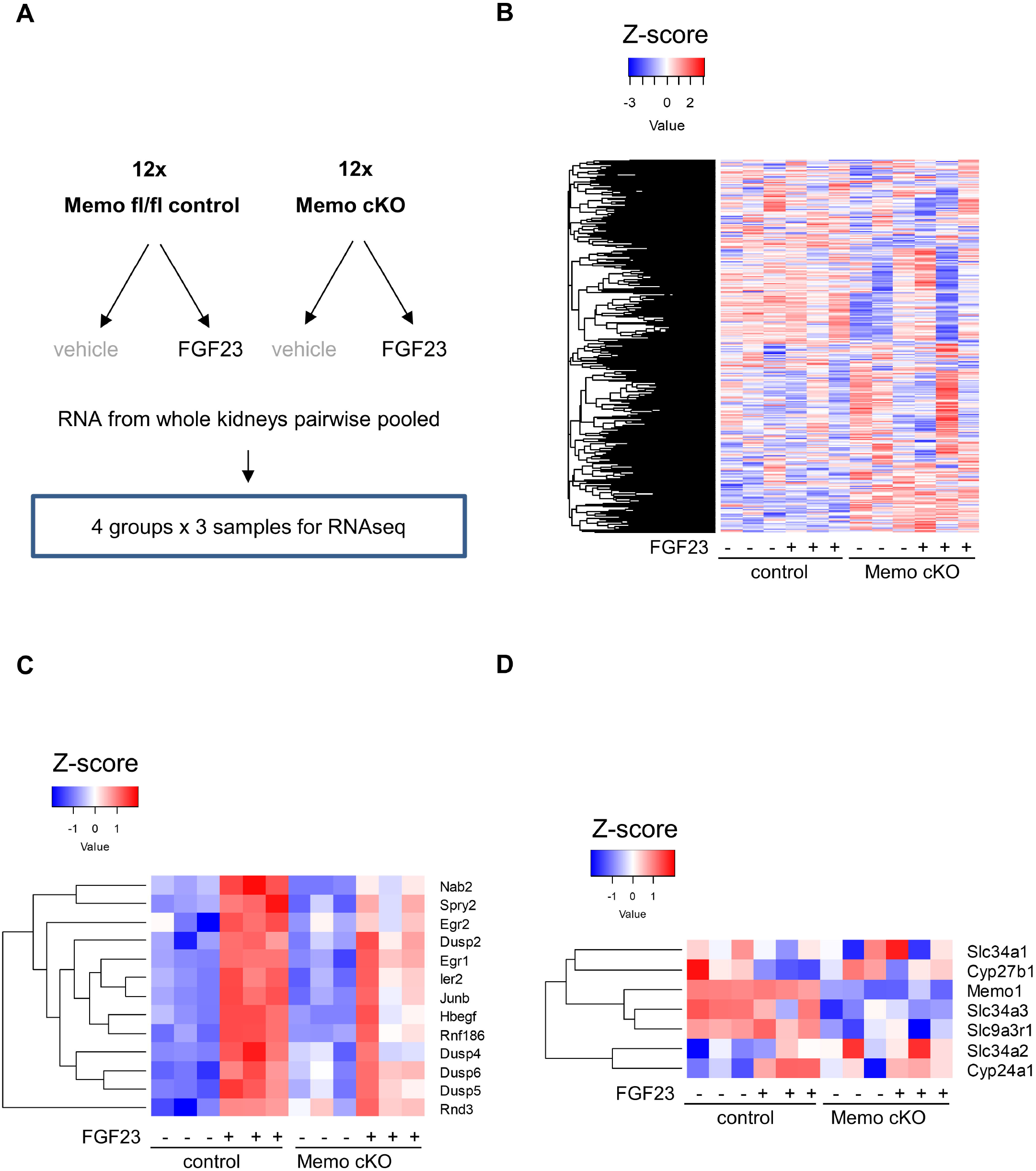
Renal RNAseq analysis shows global transcriptomic differences between Memo cKO and controls. Experimental conditions and set-up with two animals pooled per sequencing library as illustrated in (A). A heatmap of the global transcriptome shows subtle genotype differences between Memo cKO and controls (B). The 13 differently expressed transcripts between FGF23 and vehicle in c controls (C) and selected transcripts encoding proteins relevant for FGF23 signaling (D) are shown for all experimental groups.

### FGF23-FGFR driven renal PTP activity is present in controls but not in Memo-deficient animals

Next we aimed to determine if redox protein Memo affects renal FGF23 signaling via PTPs. Redox protein Memo is required for inducible NOX activity (MacDonald et al. 2014). NOXs are required during cell signaling to regulate PTPs (Lee et al. 2007), which constitutively control kinase cascades such as the ERK family. PTPs are regulated by cysteine oxidation in defined spatial areas of the cell (Miki and Funato 2012; Truong and Carroll 2012) and show an increase in phosphatase activity in response to low-dose oxidants, but a decrease at high-dose oxidants (Wright et al. 2009).

For these reasons, we aimed at deciphering downstream effectors of Memo’s redox function in FGF23-driven cell signaling, by modulating PTP activity. We treated Memo cKO and control mice with the global PTP inhibitor sodium orthovanadate 1h before applying a 1h treatment with FGF23 or vehicle. In the animals treated with PTP inhibitor, we found a diminished PTP activity in the whole kidney in sodium orthovanadate treated (Supplemental Figure 6A, right panel) compared to the control NaCl 0.9% treated animals (Supplemental Figure 6A, left panel). However, upon partial PTP inhibition by orthovanadate we discovered a relative increase in PTP activity induced by FGF23 treatment in the kidney of control animals. (Supplemental Figure 6A, right panel). This effect was blunted in Memo cKO mice. *Memo1* expression was consistently diminished in Memo cKO mice in this experimental setting, confirming the validity of the recombination in these experimental animals (Supplemental Figure 6B). To summarize, the effect of FGF23 on renal PTP activity was genotype-dependent and much stronger and less variable in animals of control genotype, yet only visible when PTPs were repressed through pharmacological inhibition.

### Oxidative modifications of the renal proteome during FGF23 signaling

We hypothesized that Memo’s redox function could disturb intracellular redox homeostasis during FGF23-driven cellular signaling. To test this, we used redox proteomics with oxidation-sensitive fluorescent probes to determine the global abundance of thiols and carbonyls as intermediates of oxidative reactions in kidney of FGF23 or vehicle-treated Memo cKO and controls. Total cellular protein thiol levels, assessed by IAF, were comparable between vehicle-treated control and Memo cKO mice (Figure 3A). However, IAF signals were increased to a higher level in FGF23-treated Memo cKO than controls, 2.54-fold and 2.17-fold respectively. These results indicate significant differences in FGF23-driven global cysteine oxidation between genotypes (Figure 3A). Total protein carbonyl content increased 1.95-fold in FGF23 treated controls but was not significantly changed in Memo cKO (Figure 3B), showing that Memo cKO kidneys abnormally process reactive intermediates (e.g. by peroxidation leading to carbonyls), resulting in excess protein thiolation in kidneys of Memo cKO. Overall, we observed distinct genotype-dependent dynamic changes in intracellular redox states during FGF23-driven signaling.

**Figure 3.**
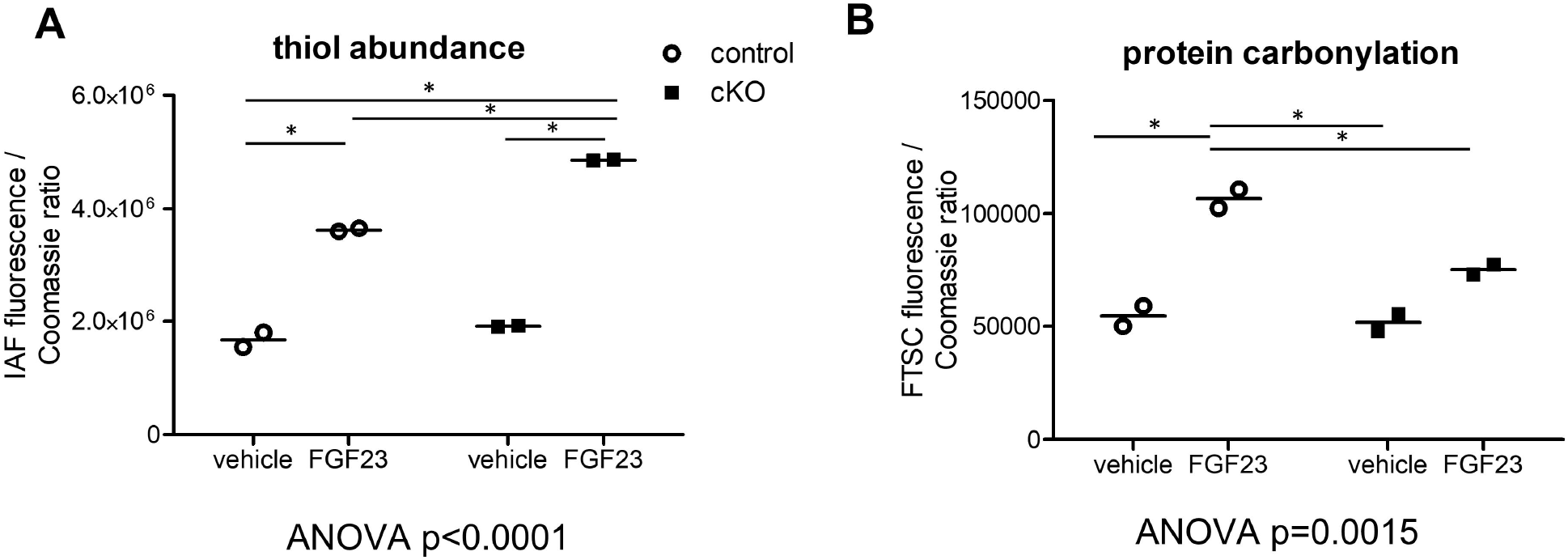
FGF23-driven change in global redox proteome in the kidney. Quantification of the redox-sensitive thiol conjugate 5-iodoacetaminofluorescein (IAF) revealed that in response to FGF23 there was a moderate 2.17-fold increase in protein thiol content FGF23 in control mice, but a 2.54-fold increase in Memo cKO (A). Global protein carbonylation measured by fluorescein-5-thiosemicarbazide (FTSC) was increased 1.95-fold by FGF23 in controls, but only 1.45-fold in Memo cKO (B). Each data point represents means of 3 technical replicates of pooled kidney halves of 2 animals (n = 2×2 per condition). Measured intensities were normalized by Coomassie stained total protein. Data were analyzed by ANOVA with Bonferroni post-tests across all columns, and significant post-tests between all groups are indicated (*).FGF23, fibroblast growth factor 23.

### A redox proteomics screen reveals interactions between Memo and the family of Rho-type small GTPases in the kidney

To identify the molecular identities of the major proteins with altered oxidation states, we reanalyzed the kidney samples shown in Figure 3A-B using 2DGE with separation by isoelectric point and mass. Subsequently, we used software-assisted determination of altered protein spots. Supplemental figure 7 shows 2DGE analyses of protein spots of control animals, where Coomassie, IAF or FTSC signals differed between genotypes. Table 1 shows the corresponding protein identities as determined by MALDI-TOF/MS. We found a diminished glutathione (GSH)-synthetase thiol signal and increased Selenium-binding protein 1 in Memo cKO. In addition to further differentially oxidized proteins associated with cytoskeleton and carbohydrate metabolism, we found one cell signaling-related protein, the Rho-GDP dissociation inhibitor 1 (Rho-GDI1), a Rho-GTPases chaperone protein. The renal thiols of Rho-GDI1 showed a 1.69-fold increase in FGF23-treated Memo cKO compared to controls (Table 1 spot no 378). To summarize, a redox-sensitive investigation of the proteome revealed an altered cysteine oxidation state of the Rho-GTPase interacting protein Rho-GDI1 between Memo cKO and controls.

We next aimed to determine the biochemical significance of this finding. We experimentally tested the possibility that Memo and Rho-GDI1 can functionally interact with each other. To this end, we analyzed whether recombinant Memo can alter the oxidation status of cysteine 79, the only cysteine in Rho-GDI1, which could potentially affect its function. To this end, we quantified posttranslational modifications of Rho-GDI1 Cys79 in native peptides and using sulfhydryl-specific iodoTMT in a cell-free environment (Figure 4A). Memo was pre-loaded with CuCl_2,_ the putative cofactor copper, followed by dialysis to remove of free copper, as previously described (16). We found that the reversible oxidation of Rho-GDI1 Cys79 to sulfenate was negligible, but irreversible dioxidation to sulfinyl groups (SO2H) and to some extent trioxidation to sulfonyl groups (SO3H) at Cys79 of Rho-GDI1 was detected (Figure 4B). H2O2 served as a positive control for irreversible oxidation. These findings suggest that Memo directly oxidizes Rho-GDI1 in vitro.

**Figure 4.**
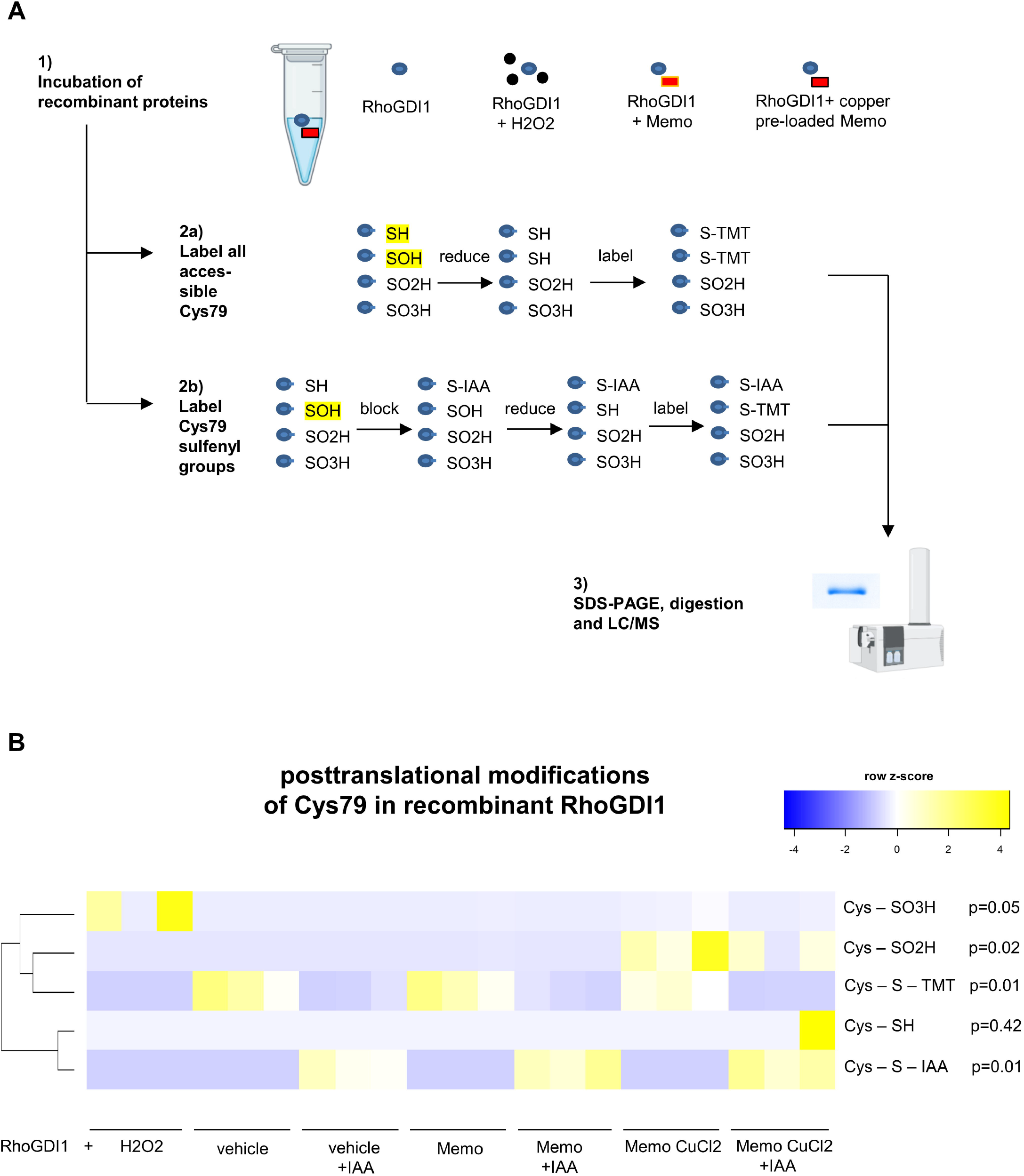
Oxidative interaction between recombinant Memo and Rho-GDI1 in cell-free condition. A shows the experimental workflow for modification analyses of Cys79 in recombinant Rho-GDI1, after incubation with or without copper pre-loaded Memo or H2O2. In brief, irreversibly dioxidized (SO2H) and trioxidized (SO3H) were directly detected by MALDI-TOF/MS, and reversible monooxidation (SOH) peptides were detected using a iodoTMT tandem mass-spectroscopy tag (TMT) after blocking accessible cysteines by iodoacetamide (IAA); total accessible cysteines were detected without IAA blocking. Incubation of RhoGDI1 with H2O2 as a positive control caused trioxidation (B, first row), and incubation with copper-preloaded and dialyzed Memo caused dioxidation and to a lesser extent trioxidation in RhoGDI1 Cys79 (B, blue rectangle). Memo or copper-preloaded Memo addition did not alter the TMT label intensity (B, 3^rd^ row and blue rectangle). Only one sample had some remaining free thiols (B, 4^th^ row), and in all IAA-blocked samples, the IAA alkylation was present (B, 5^th^ row). Signal intensities in B are normalized for total number intensity of detected peptides, and displayed as z-scores of original non-transformed data. or Kruskal-Wallis test with Dunn’s multiple testing correction (B). *, p< 0.05 in post-tests. n=3 independent experiments per condition.

**Figure 5.**
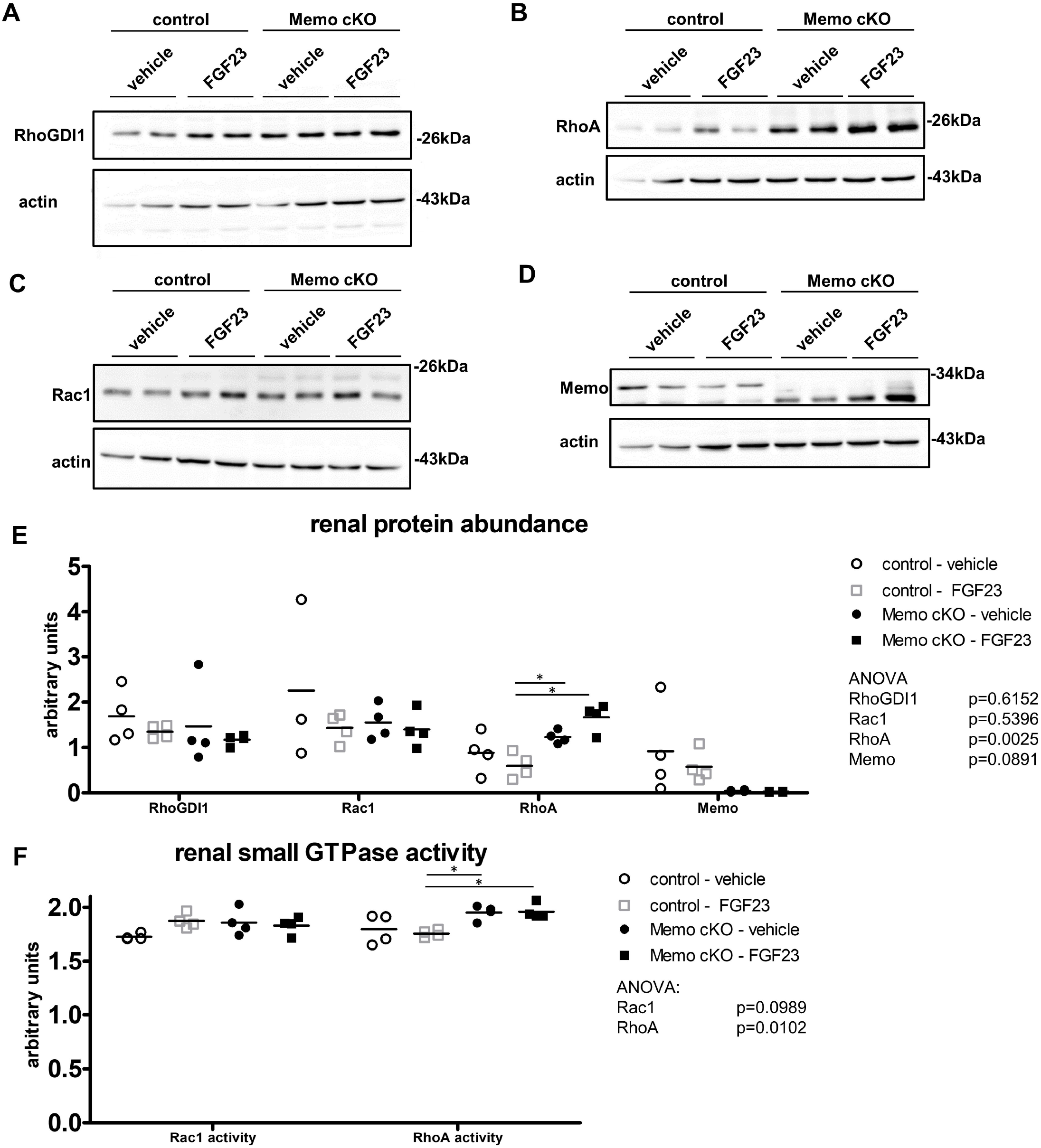
Malfunctioning Rho-GTPase network in kidneys from Memo cKO (cKO) mice. Western blots of whole kidneys are shown for indicated proteins (A-D) and their respective densitometric quantification against actin as a loading control in. Renal Rac1 and RhoA activity were measured by G-LISA assays (F). Data in E-F were analyzed by ANOVA with Bonferroni correction (6 post-tests each). *, post-test p<0.05. n=4 per condition in A-F. n=4 per condition (A-F); except n=3 in E (Rac1: control vehicle).

Next, we aimed at delineating whether Rho-GDI1 is directly required in the early signaling cascades of FGF23-driven signaling, or if it rather plays a less direct role in this process by influencing other mediators. We performed genetic manipulations of Rho-GDI1 in a cellular system of HEK293 cells that stably express Klotho (HEK293-KL) (Supplemental figure 8) (Diener et al. 2015). Rho-GDI1 is endogenously expressed in HEK293 cells (Boulter et al. 2010). Using two different siRNAs targeting Rho-GDI1, we knocked down Rho-GDI1 to 1% or 30% of mock-transfected cells (Supplemental figure 9A-B and 10A-B). Next, we assessed the cellular signaling responses to FGF23 at two different time points. There was no difference in ERK phosphorylation in response to FGF23 stimulation between Rho-GDI1 knockdown and control cells (Supplemental figure 9C-D and 10C-D). Conversely, we tested if increased abundance of Rho-GDI1 affects FGF23 signaling. We transiently overexpressed Rho-GDI1 in (Supplemental figure 11A). Again, there was no difference in ERK phosphorylation in response to FGF23 stimulation between Rho-GDI1 overexpressing and mock-transfected cells (Supplemental figure 11B-C). Overall, knock down or overexpression of Rho-GDI1 in HEK293-KL cells did not alter ERK phosphorylation levels in response to FGF23.

Rho-GDI1 is a chaperone protein and regulator of small Rho-GTPases such as RhoA and Rac1, two proteins that are particularly important for cell signaling including FGFR signaling (13). Rac1 is required for NOX-mediated inactivation of PTPs via cysteine oxidation to promote RTK signaling (Wright et al. 2009; Miki and Funato 2012; Truong and Carroll 2012). We have shown in Figure 4 that Memo directly oxidizes Rho-GDI1. Moreover, Memo can directly oxidize RhoA (MacDonald et al. 2014). Next, we investigated if overall protein level of Rho-GDI1, Rac1 and RhoA are altered in Memo cKO kidneys.

Rho-GDI1 abundance is an important regulator of Rho-GTPase activity overall (Boulter et al. 2010). Therefore, we tested the overall abundance of Rho-GDI1 in the kidney of Memo cKO and control animals. Rho-GDI1 abundance was comparable between genotypes and FGF23 treatment status (Figure 5A, quantification in Figure 5E). With total Rho-GDI1 unchanged, the decreased Rho-GDI1 thiols in FGF23-treated control mice would correspond to a transient oxidation of Rho-GDI1 Cys79 to sulfenate in order to liberate and activate a bound Rho-GTPase. A similar observation was previously reported for the RhoA – Rho-GDI1 complex (Kim et al. 2017). Memo is required for growth factor-driven RhoA relocalization to the membrane (Zaoui et al. 2008), and strikingly, RhoA protein abundance and activity were increased in the kidney of Memo cKO mice in both vehicle and FGF23 treated cohorts (Figure 5B, 5E-5F). The abundance of the Rho-GTPase Rac1 was unchanged in all conditions (Figure 5C, quantification in Figure 5E), however, FGF23 tended to increase Rac1 activity by 10% in the whole kidney of FGF23 treated control mice (Figure 5F), similar to what has been reported for Rac1 activity in response to FGF2 treatment in cell culture (Shin et al. 2006; Barrios and Wieder 2009). Importantly, there was no such trend towards increased Rac1 activity in FGF23 treated Memo cKO animals. Renal Memo protein was diminished in Memo cKO as expected (Figure 5D-E). To conclude, in addition to the increased Rho-GDI1 thiol signal, we found a major increase in abundance and activity of the renal Rho-GDI1 interacting protein RhoA due to *Memo1* loss-of-function.

## Discussion

Memo is an intracellular moonlighting protein involved in a plethora of cellular processes (Schotanus and Van Otterloo 2020). These include cell cycle regulation, glucose metabolism and cytoskeletal dynamics (Zaoui et al. 2008; Ding et al. 2020; Schotanus and Van Otterloo 2020). Memo promotes cellular responses to phosphorylation of different RTKs (Marone et al. 2004), and Memo associates with intracellular adaptor proteins of the RTK signalosome (Marone et al. 2004; Sorokin and Chen 2013; Haenzi et al. 2014). In a mouse model of postnatal inducible Memo deficiency in the whole body, we have previously reported a unique syndrome of accelerated aging, insulin hypersensitivity, bone disease, elevated FGF23 and calcium concentrations in serum and increased renal phosphate cotransporter NaPi2a expression that recapitulates many features of *Klotho* or *Fgf23* loss of function models (Haenzi et al. 2014; Moor et al. 2018; Moor et al. 2020). Here, we set out to discover mechanisms underlying Memo’s role in RTK signaling using FGF23-driven renal signaling as an example.

In Memo-deficient mice injected with FGF23, we demonstrated Memo’s importance by showing blunted early renal ERK phosphorylation events and transcriptional responses upon FGF23 treatment. The differently expressed genes after FGF23 treatment in control mice including DUSP and early response proteins closely resembled the findings from Ni et al. who used a higher dosage and the identical treatment duration (Ni et al. 2021), and these responses were diminished in Memo cKO. The plethora of signaling pathways affected by *Memo1* loss-of-function includes not only FGFR signaling (Schotanus and Van Otterloo 2020). The observed FGF23 signaling deficit does likely not result from alterations in FGFR subtype composition or Klotho protein in Memo cKO animals. Klotho protein abundance was also preserved in the animals studied.

Moreover, Memo protein has no kinase or phosphatase activity (Qiu et al. 2008) that could directly influence MAPK signaling. Therefore, we asked whether Memo’s redox function might be involved in signal regulation. Indeed, RTK-dependent signal propagation through kinases requires oxidative conformational change of PTPs that releases their inhibitory effects (Miki and Funato 2012; Truong and Carroll 2012) (see Figure 6). Moreover, by partially repressing PTPs to a submaximal state of activity using a PTP inhibitor, we found that PTP activity can be increased by FGF23. This is, to our knowledge, the first evidence that FGF23 activates this negative feedback loop in promoting PTP activity in the kidney that could then dephosphorylate FGFRs. We found FGF23-driven PTP activation in kidneys of controls but not Memo-deficient animals which presented a more variable phenotype, potentially due to disturbed cellular redox homeostasis. This prompted us to further investigate the oxidative changes occurring during FGF23 signaling in these animals.

**Figure 6.**
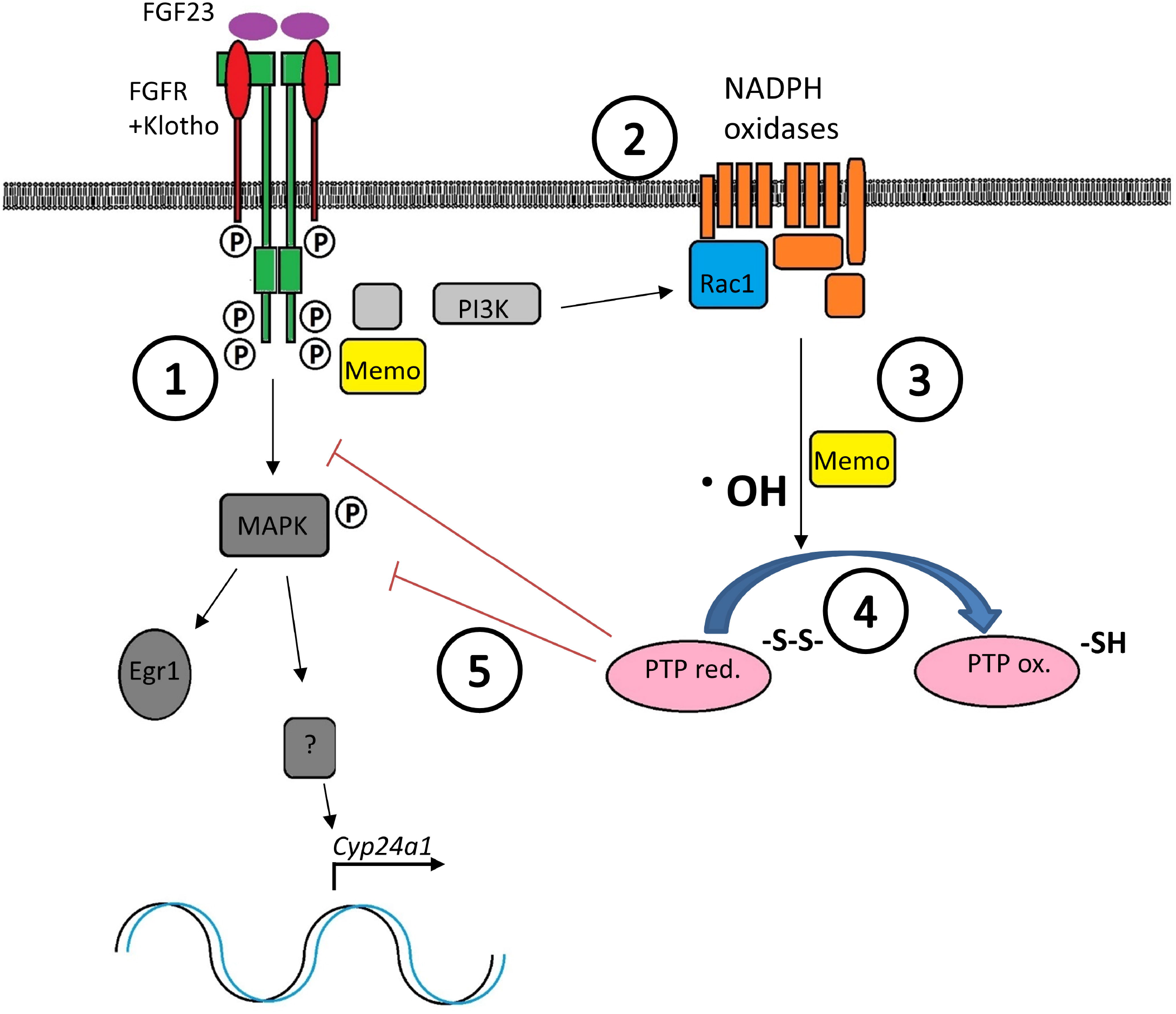
Model illustrating Memo-dependent FGF23-driven cell signaling in the kidney. (1) FGF23 induces phosphorylation and dimerization of the FGFR and recruits intracellular adaptor proteins including FRS2, GAB1, GRB2, and Memo (Ref.: Haenzi H et al., FASEB J 2014). (2) RAS-dependent PI3K activity induces Rac1 GTPase activation and formation of multiprotein assemblies forming NADPH oxidases (Ref.: Chiarugi P, TRENDS Biochem Sci 2003). (3) Memo promotes the intracellular ROS generation by the NADPH oxidases (Ref.: Macdonald G et al., Sci Signal 2014). (4) ROS such as ·^•^**OH** then oxidixe Cysteines in PTPs, causing a conformational change and inactivation of phosphatase activity (Ref.: Chiarugi P, TRENDS Biochem Sci 2003). (5) Decreased phosphatase activity of PTPs promotes the kinase activity of MAPK pathway enzymes, allowing nuclear translocation of transcription factors that drivetarget gene expression of *Cyp24a1*. PI3K, phopshatidyl-inositol-3-phosphate kinase. PTP, protein phosphotyrosyl-phosphatase. S-S reduced cysteine sulfur. SH, thiol. ROS, reactive oxygen species. ·OH, hydroxyl radical as a proxy for reactive oxygen species. Parts of the illustration are adapted from (Ref.: Chiarugi P, TRENDS Biochem Sci 2003).

We performed redox proteomics and found a change in the Rho-GTPases chaperone protein Rho-GDI1. The renal thiols of Rho-GDI1 were increased in FGF23-treated Memo cKO compared to controls. This finding was validated and further analyzed by biochemical studies with selected recombinant proteins, followed by studies of kidney lysates. Our results show that the Rho-GTPase network in the kidney is altered in Memo’s absence.

The interaction between Memo and Rho-GTPases results in a differential regulation of multiprotein subunit recruitment of NADPH oxidases in other model systems (MacDonald et al. 2014; Ewald et al. 2017). This has implications for the well-characterized intracellular redox function of Rac1 needed for NADPH oxidase assembly (Flinder et al. 2011). Importantly, both RhoA and Rac1 regulate RTK endocytosis, which is a cue to terminate FGFR-mediated MAPK signaling. Results from cysteine oxidation proteomic studies have recently highlighted extensive alterations in the oxidation of the Rho-GTPase subfamilies RHO and RAC in response to epidermal growth factor-driven RTK signaling (Lamaze et al. 1996; Bryant et al. 2005; Grassart et al. 2008; Behring et al. 2020). Both of these Rho-GTPases as well as their chaperone protein Rho-GDI1 are expressed in the renal tubular segments relevant for FGF23-driven signaling (Gee et al. 2013; Steichen et al. 2021).

Next, we found a diminished GSH-synthetase thiol signal and increased Selenium-binding protein 1 in Memo cKO point. This points towards a relevance of Memo for selenium-and/or GSH-containing cellular antioxidative protection systems (Zhang et al. 2020) to potentially prevent oxidative damage and excessive uncontrolled protein thiol formation during RTK signaling. This is supported by a recent redox biochemistry study by Zhang et al. who found that Memo1 Protects copper from initiating redox cycling in vitro (Zhang et al. 2022).

This study contains some limitations. First, the phenotype of Memo deficiency has temporal dynamics and was assessed only at a single time point before onset of chronic kidney disease (Haenzi et al. 2014; Moor et al. 2018). As done in the past (Ni et al. 2021), we studied acute signaling events with a focus on selected FGF23-dependent intracellular pathways. Later time points after FGF23 treatment would be required to induce the physiological effect of downregulating NaPi2a phosphate transport protein to promote phosphaturia, as reported from experiments with isolated renal tubules (Andrukhova et al. 2012). Most findings using the Memo-floxed allele showed a degree of variability, similarly as reported earlier (Haenzi et al. 2014). A potential explanation for this could be a truncated Memo1 protein lacking exon 2 that is variably degraded and exerting some partial function in Memo1 mutant animals. Next, we could not capture the precise role for Rho-type GTPases in FGF23 signaling in this setting. GDP/GTP exchange and GTPase activities of Rho-type GTPases underlie posttranslational regulation mechanisms such as phosphorylation, cysteine oxidation or compartmental trafficking that should be investigated in future biochemical studies of RTK signaling (Hobbs et al. 2014; Behring et al. 2020). Finally, the PTPs that modulate FGF23 signaling may include several redundant enzymes that are not easily distinguished *in vivo* or captured using the chosen gel-based proteomics approach but could be further addressed *ex vivo* using purified complexes. In brief, further study of the moonlighting protein Memo and other redox proteins may provide many opportunities to discover currently elusive regulators of RTK signaling.

To conclude, the presented data reveal an exciting biological relevance for Memo at the intersection of RTK signaling, redox homeostasis and intracellular feedback regulation by PTPs after stimulation with FGF23. This member of the FGF family is still incompletely understood as a hormone, biomarker, and drug target in bone, kidney and cardiovascular disease.

## List of abbreviations

2DGE: 2-dimensional gel electrophoresis
ANOVA: analysis of variance
ARHGDIA: Rho GDP Dissociation Inhibitor Alpha
BSA: bovine serum albumin
CKD: chronic kidney disease
cKO: conditional knockout
Cyp24a1: Cytochrome P450 Family 24 Subfamily A Member 1
Cyp27b2: Cytochrome P450 Family 27 Subfamily B Member 2
Cys: cysteine
DTT: dithiothreitol
DUSP: dual-specificity phosphatase
EDTA: ethylenediaminetetraacetic acid
Egr1: Early growth response 1
ERK: extracellular signal-regulated kinase
FGF: fibroblast growth factor
FGFR: fibroblast growth factor receptor
fl/fl: floxed
FRS2: FGFR substrate protein
FTSC: fluorescein-5-thiosemicarbazide
GAPDH: glyceraldehyde-3-phosphate dehydrogenase
GDP: guanosine diphosphate
GTP: guanosine triphosphate
Hbegf: heparin-binding epidermal growth factor-like growth factor
HEK293: human embryonic kidney cells 293
HEK293-KL: HEK293 stably expressing Klotho
IAF: 5-iodoacetamido fluorescein
IEF: isoelectric focusing
Il17rd: Interleukin 17 Receptor D
IPG: immobilized pH gradient
LC: liquid chromatography
MALDI-TOF/MS: Matrix Assisted Laser Desorption/Ionization coupled to time-of-flight mass spectrometry
MAPK: mitogen-activated protein kinase
NaPi2a: sodium-dependent phosphate transport protein 2A
Memo1: Mediator of Cell Motility 1
NADPH: nicotinamide adenine dinucleotide phosphate
NOX: NADPH oxidase
NP-40: Nonidet P-40
PBS: phosphate buffered saline
PCR: polymerase chain reaction
pERK: phosphorylated ERK
PMSF: phenylmethylsulfonyl fluoride
PTP: protein tyrosyl phosphatase
PVDF: polyvinylidene fluoride or polyvinylidene difluoride
Rho-GDI1: Rho-GDP dissociation inhibitor 1
Rac1: Rac Family Small GTPase 1
RCM: reduction and carboxymethylation
RIPA: radioimmunoprecipitation assay
RNAseq: RNA sequencing
RT: reverse transcriptase
RTK: receptor tyrosine kinase
SDS-PAGE: sodium dodecyl sulfate–polyacrylamide gel electrophoresis
SEF: similar expression to fibroblast growth factor
siRNA: small interfering RNA
TBST: Tris-buffered saline supplemented with Tween
tERK: total ERK
TMT: tandem mass tag

## Acknowledgements

The authors wish to thank Daniel Bardy, Laboratoire Central de Chimie, Lausanne University Hospital for the analysis of mouse serum samples; Craig Pattinson and Daniel Stead, University of Aberdeen for protein identification and Ivan Nalvarte, Karolinska Institutet, Stockholm, for protocol and reagent sharing. The authors thank members of the Proteomics Core Facility of the University of Bern for support with analyses, and Sandra Calderon and Leonore Wigger at Genomic Technologies Facility, University of Lausanne for RNAseq data acquisition and analysis. The authors thank Bettina Lorenz-Depiereux, Helmholtz Zentrum München for providing the HEK293-Klotho cell line. The authors thank Uyen Huynh-Do and Stefan Rudloff, Department of Nephrology and Hypertension, University Hospital Bern for helpful discussions and acknowledge infrastructure support by the Department of Nephrology and Hypertension, University Hospital Bern.

## Author contributions

MBM and BH conceived the study. KB, SKR, SB, BH, DS, OB and MBM participated in experimental design. KB, SKR, SB, FD, AP, YZ, and MBM performed experiments. KB, SKR, DS, NEH, OB and MBM participated in data analysis and interpretation. DS and NEH provided crucial laboratory materials. OB and MBM acquired funding for the research. OB and MBM supervised the research. MBM wrote the manuscript. All authors critically read and commented on the manuscript and agreed to manuscript submission.

