## Supplemental Figures for "Redox protein Memo1 coordinates FGF23-driven signaling and small Rho-GTPases in the mouse kidney"

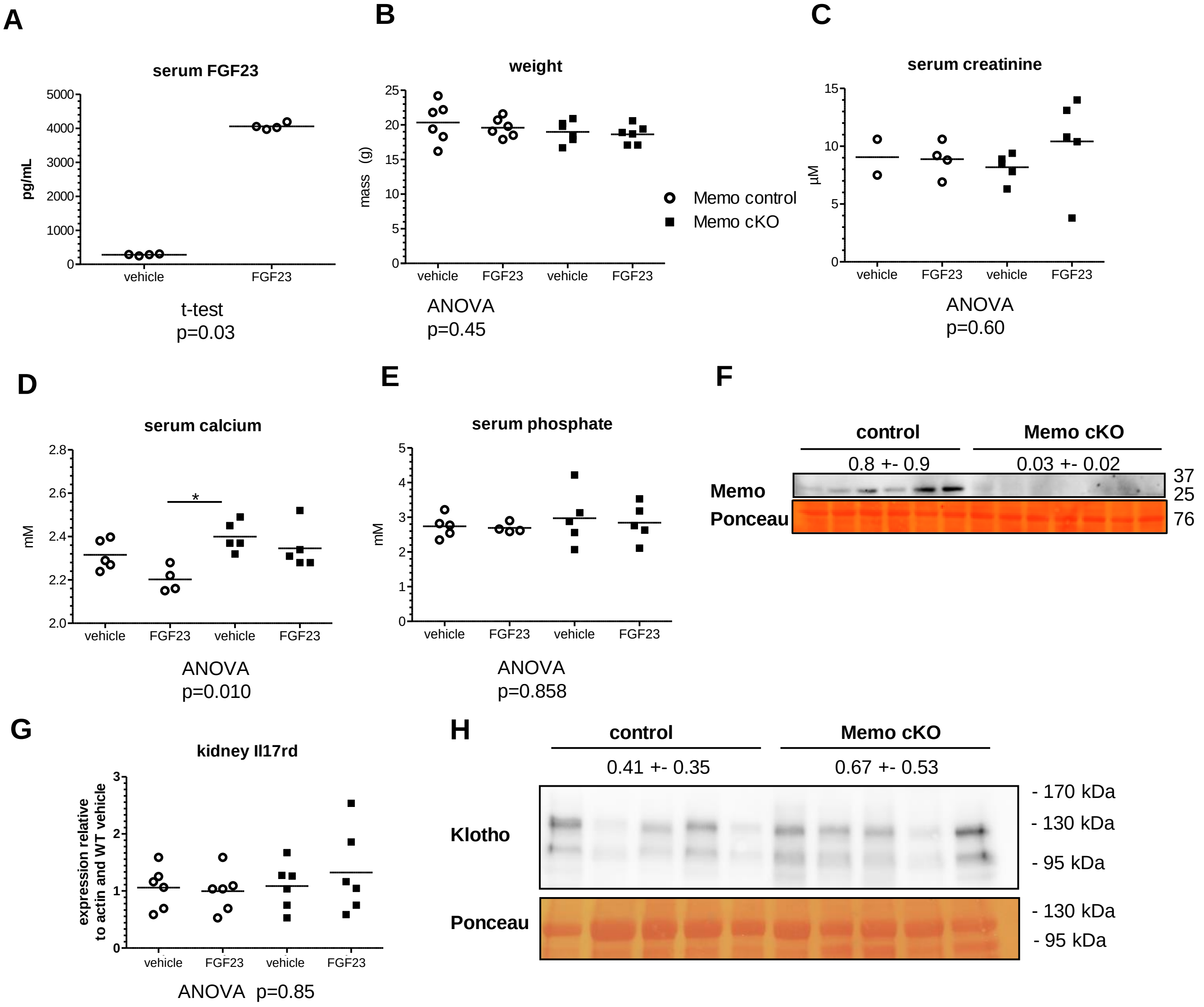


**Supplemental figure 1. Characteristics of control and Memo cKO mice treated with FGF23 or vehicle.** Mice showed normal body weight (B). Serum creatinine was constant across all experimental conditions (C). Serum calcium concentrations varied between conditions, with slightly elevated levels in vehicle-treated Memo cKO (D). Serum phosphate concentrations were comparable across all experimental groups (E). Ablation of Memo protein was verified in kidney by Western blot (F); densitometric quantification indicated (mean+-standard deviation), t-test p=0.046). Target gene responses of FGFR regulator SEF encoded by *Il17rd* was assessed by qPCR (G). FGF23 co-receptor Klotho protein levels at 130 kDa (H) were comparable in the kidney across experimental groups; quantification indicated of Klotho bands at 130kDa (t-test; p=0.38). Sample size is indicated by the number of data points in scatterplots (B: 6 per group, C: 2 to 5 per group; D: 4 to 5 per group; E: 4 to 5 per group; F: 6 controls and 7 cKO;  G: 6 per group; H: 5 per group). Analysis by ANOVA (B-E, G) followed by Bonferroni's or Dunn's post-test across all experimental groups with * p<0.05 in post-tests.

A B

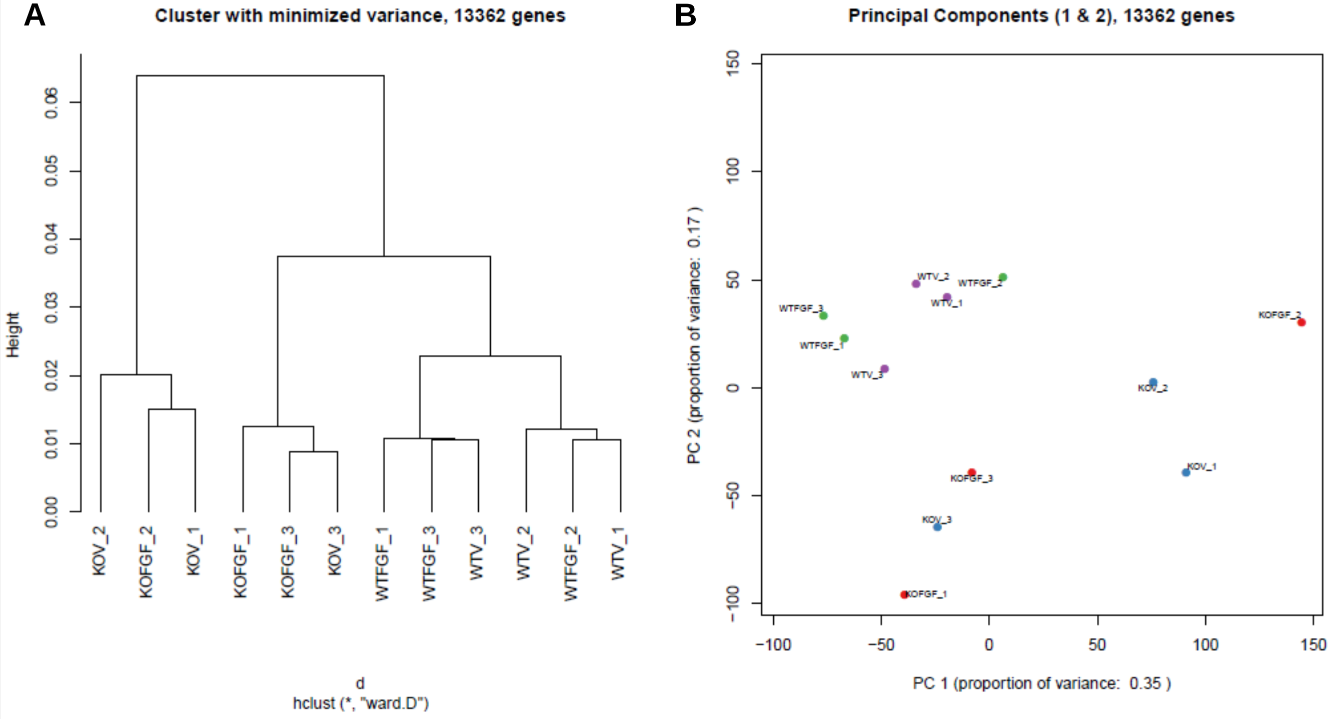


C

| Comparison | P adj. < 0.05 | | | P adj. < 0.05 and fold change > 2 | | |
|---|---|---|---|---|---|---|
| | ALL | UP | DOWN | ALL | UP | DOWN |
| List1 KOFGF_vs_KOV | 1 | 1 | 0 | 1 | 1 | 0 |
| List2 WTFGF_vs_WTV | 13 | 13 | 0 | 11 | 11 | 0 |
| List3 KOFGF_vs_WTFGF | 88 | 71 | 17 | 63 | 49 | 14 |
| List4 KOV_vs_WTV | 85 | 61 | 24 | 56 | 43 | 13 |
| List5 (KOFGF_vs_KOV)_vs_(WTFGF_vs_WTV) | 1 | 1 | 0 | 1 | 1 | 0 |

**Supplemental figure 2. Global analysis of 13362 protein-coding renal transcripts in control and Memo cKO mice treated with FGF23 or vehicle.** Harvesting kidneys 1h after FGF23 injection did not reveal a global impact; the only differences seen in unsupervised clustering (A) and principal component analysis (B) were in the genotype. C shows the number of transcripts affected by the treatment or genotype, where "KOFGF-KOV„ is the treatment effect in knockout, "WTFGF-WTV" is the treatment effect in wild type, "KOFGF-WTFGF„ is the knockout effect in treated mice, "KOV-WTV" is the Memo cKO effect in vehicle-treated mice and "(KOFGF-KOV)-(WTFGF-WTV)" is the interaction of genotype and treatment across all 4 experimental groups. adj., adjusted.

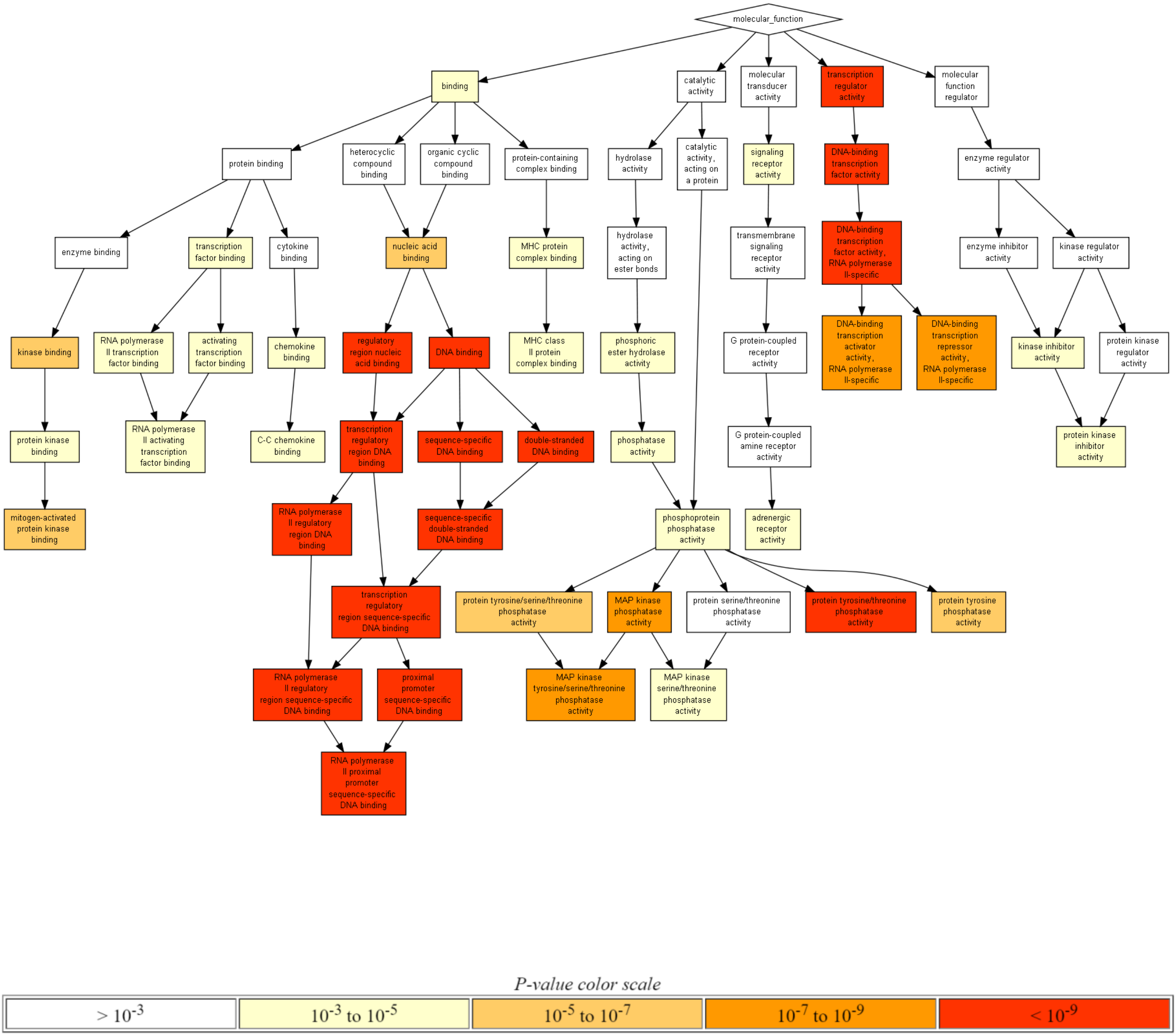


**Supplemental figure 3. Gene ontology analysis of FGF23 transcriptional effects in kidney of control mice.** MAP, mitogen-activated protein. False discovery rate-adjusted q values are indicated in Supplemental table 2.

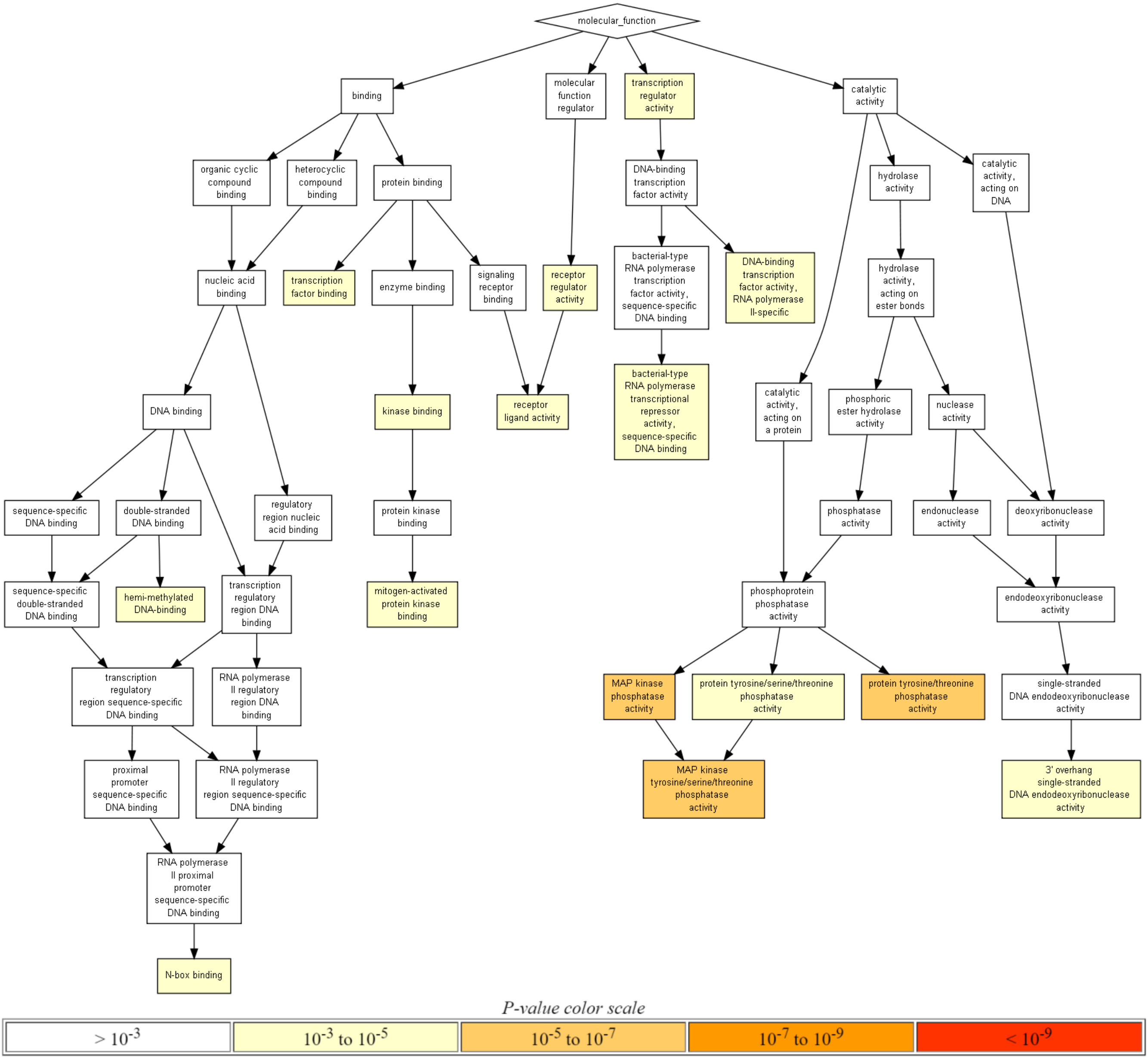


**Supplemental figure 4. Gene ontology analysis of FGF23 transcriptional effects in kidney of Memo cKO mice.** MAP, mitogen-activated protein. False discovery rate-adjusted q values are indicated in Supplemental table 1.

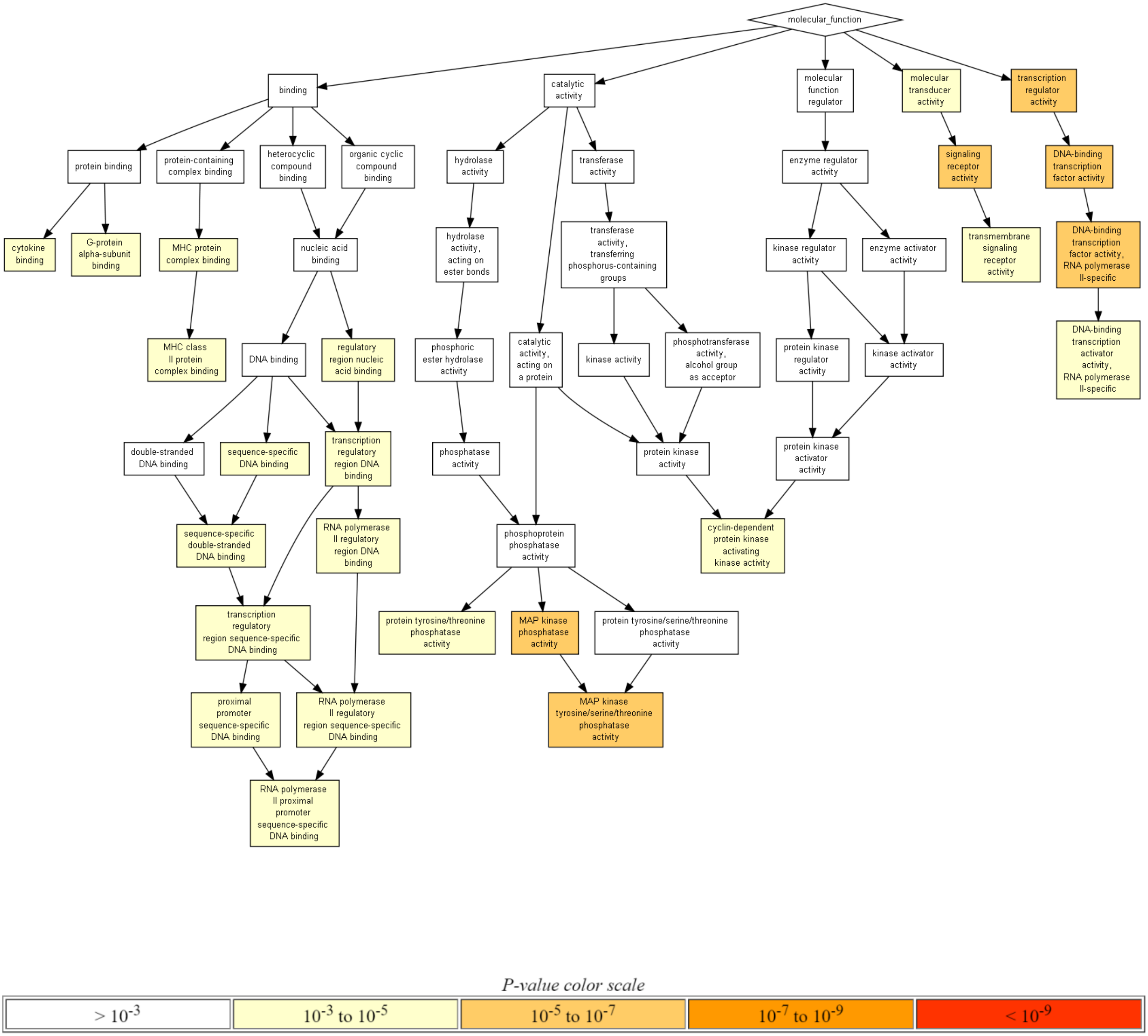


**Supplemental figure 5. Gene ontology analysis of renal transcriptomic interaction between FGF23 treatment effect and genotype effect.** MAP, mitogen-activated protein. False discovery rate-adjusted q values are indicated in Supplemental table 5.

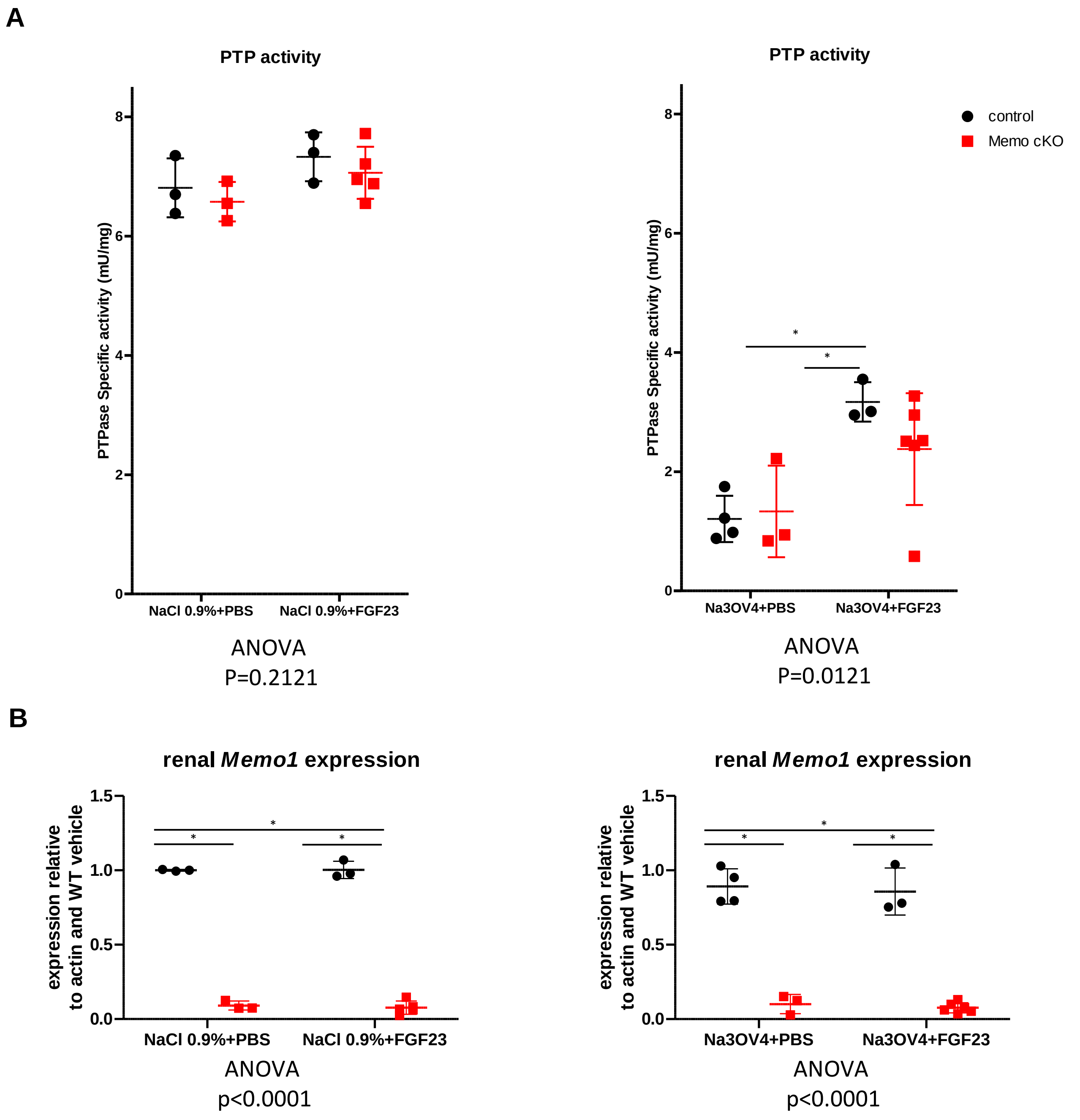


**Supplemental figure 6. FGF23-driven protein phosphotyrosyl phosphatases (PTP) activity in the whole kidney after inhibition of protein phosphotyrosyl phosphatases.** Sodium orthovanadate partially inhibited PTP activity in the kidney (A, right panel) compared to vehicle (A, left panel) and allowed the detection of an FGF23-driven relative increase in PTP activity in control mice but not in Memo cKO mice (A, right panel). Memo1 gene expression was diminished in the kidney of Memo cKO in comparison to controls (B). Statistical analysis by ANOVA with Bonferroni post-tests across all columns. N=3 to 6 mice per genotype with each scatter representing one mouse. NaCl, sodium chloride. Na3OV4, sodium orthovanadate. PBS, phosphate-buffered saline. FGF23, fibroblast growth factor 23.

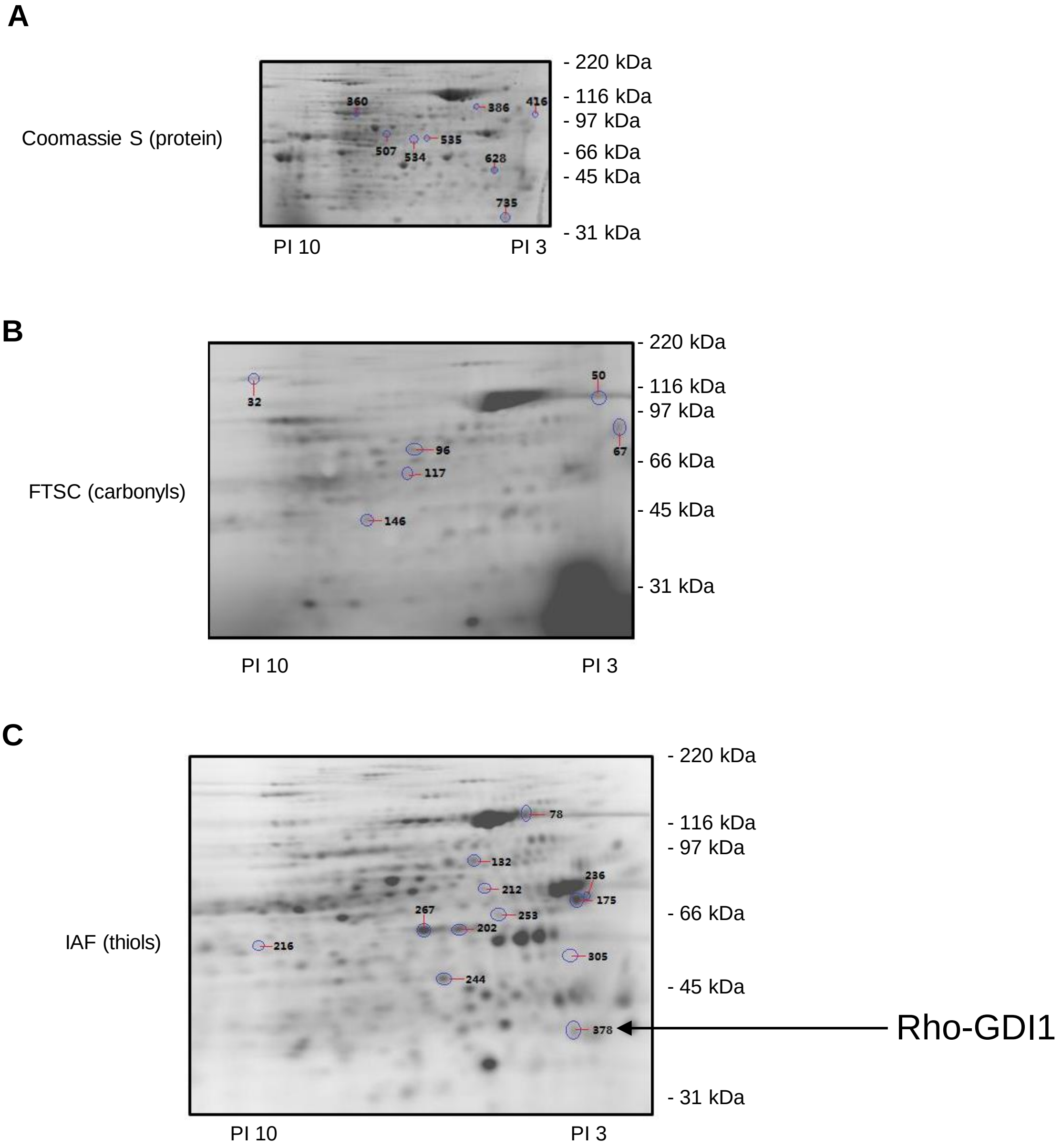


**Supplemental figure 7. Two-dimensional gel electrophoresis of kidney lysates of control mice used for protein identification after redox proteomic screens.** Samples pooled in pairs (n=2x2 per condition) separated by isoelectric point and size, followed by determination of altered protein spots in technical triplicates. Coomassie stains for total protein (A), fluorescent label FTSC for carbonyls (B) and IAF for thiols (C) are shown including the individual differently abundant protein spots of proteins presented in Table 1. IAF, 5-iodoacetaminofluorescein. FTSC, fluorescein-5-thiosemicarbazide.

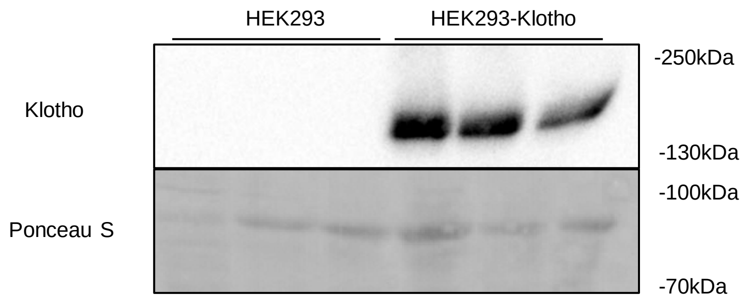


**Supplemental figure 8.** HEK293-Klotho cells stably transfected with Klotho show a detectable Klotho-specific band at 130-150 kDa, whereas non-transfected HEK293 cells do not express Klotho.

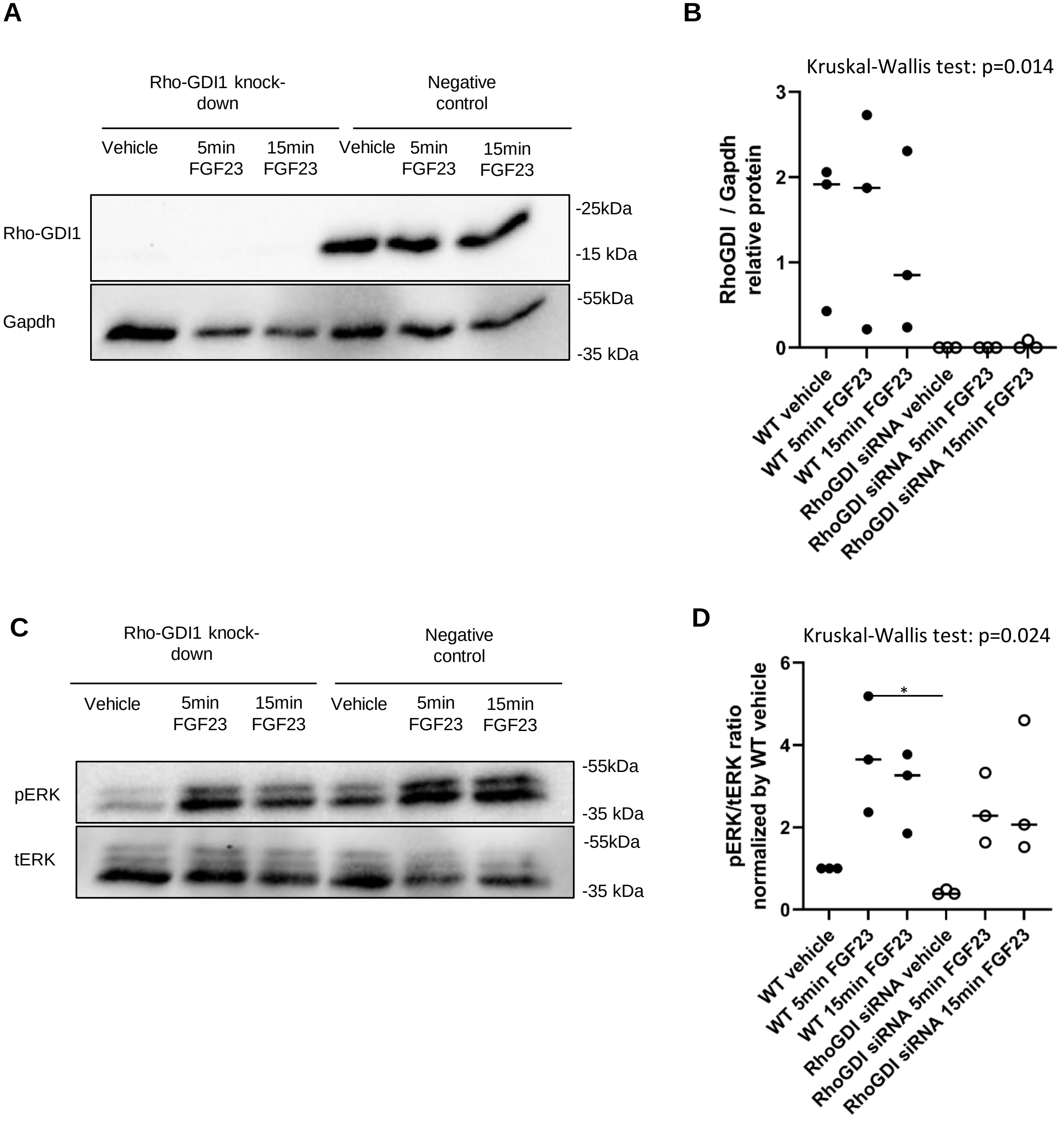


**Supplemental figure 9. siRNA knockdown (Origene) of Rho-GDI1 in Klotho-expressing cells does not alter FGF23-driven ERK phosphorylation.** In HEK293-Klotho cells stably transfected with Klotho, an siRNA targeting Rho-GDI1 from Origene led to efficient depletion of Rho-GDI1 protein compared to a non-targeting control siRNA (A), densitometric quantification in B. FGF23 caused an increase in ERK phosphorylation at the two indicated time points that was comparable between Rho-GDI1 depleted cells versus cells treated non-targeting control siRNA (C), quantification in D. Statistical analysis was obtained by Kruskal Wallis test with Dunn’s multiple testing correction performed between all groups. N=3 independent experiments per experimental condition.

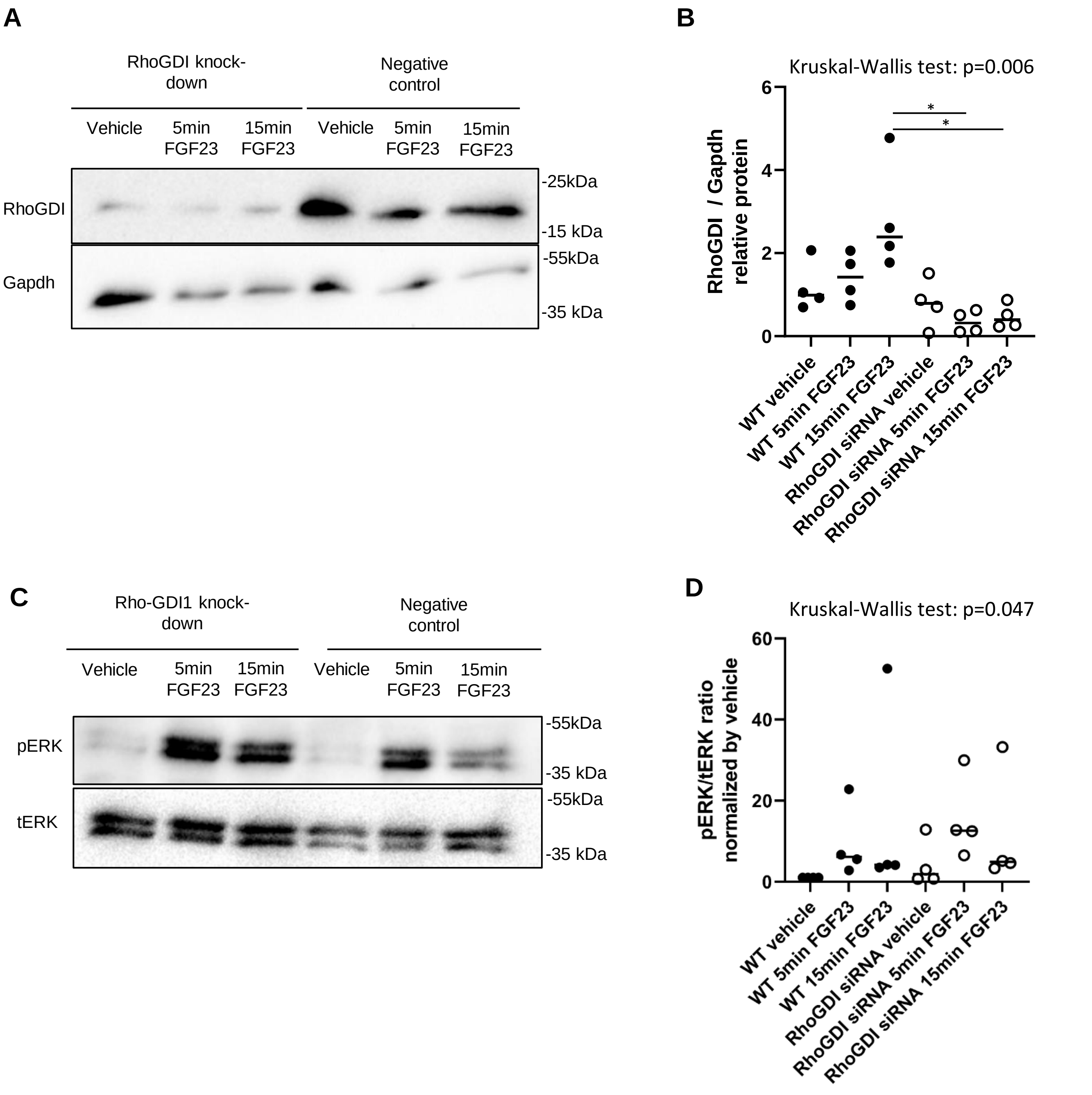


**Supplemental figure 10. siRNA knockdown (ThermoFisher) of Rho-GDI1 in Klotho-expressing cells does not alter FGF23-driven ERK phosphorylation.** In HEK293-Klotho cells stably transfected with Klotho, an siRNA targeting Rho-GDI1 from Thermo Fisher led to efficient depletion of Rho-GDI1 protein compared to a non-targeting control siRNA (A), densitometric quantification in B. FGF23 caused an increase in ERK phosphorylation at the two indicated time points that was comparable between Rho-GDI1 depleted cells versus cells treated non-targeting control siRNA (C), quantification in D. Statistical analysis was obtained by Kruskal Wallis test with Dunn's multiple testing correction performed between all groups. N=3 independent experiments per experimental condition.

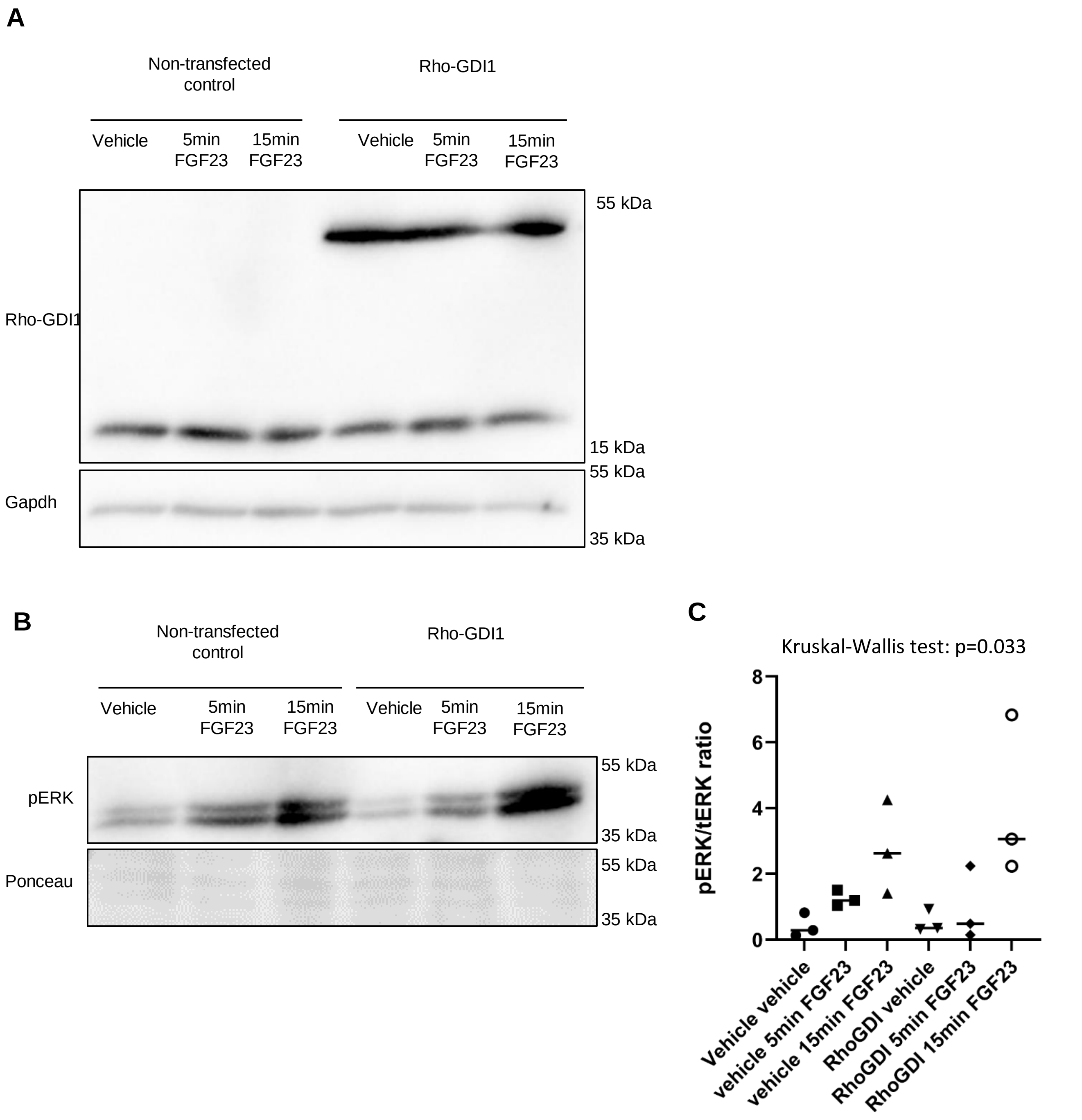


**Supplemental figure 11. Overexpression of Rho-GDI1 in Klotho-expressing cells does not alter FGF23-driven ERK phosphorylation.** In HEK293-Klotho cells stably transfected with Klotho, Rho-GDI1 was overexpressed leading to a multimeric band of around 50 kDa that was recognized by Rho-GDI1 antibody in Western blot (A). FGF23 caused an increase in ERK phosphorylation at the two indicated time points that was comparable between Rho-GDI1 overexpressing cells versus mock-transfected control cells (B), quantification in C. Statistical analysis was obtained by Kruskal Wallis test with Dunn’s multiple testing correction performed between all groups. N=3 independent experiments per experimental condition.
